# Postnatal maternal care normalizes the hypothalamic DNA methylome following prenatal bisphenol exposure

**DOI:** 10.1101/2025.10.03.680379

**Authors:** Samantha C. Lauby, Dennis C. Wylie, Hannah E. Lapp, Melissa Salazar, Amy E. Margolis, Frances A. Champagne

## Abstract

**Background:** Environmental exposures co-occurring during early life have a profound influence on neurodevelopment. Our previous work in rats suggests that postnatal maternal care modulates the effects of prenatal exposure to bisphenols, an estrogenic endocrine disrupting chemical, on offspring neurodevelopment. Elevated maternal licking/grooming and prenatal bisphenol exposure have opposing effects on estrogen receptor expression in the medial preoptic area (MPOA), which could impact expression of estrogen-responsive genes. We hypothesized that maternal licking/grooming would mitigate the effects of prenatal bisphenol exposure on estrogen receptor signaling in the developing MPOA. In addition, we hypothesized that there would be interactive effects of prenatal bisphenol exposure and maternal licking/grooming on DNA methylation nearby estrogen responsive elements.

**Results:** Our results suggest that maternal licking/grooming normalized prenatal bisphenol-induced upregulation of estrogen-related receptor gamma (*Esrrg*) expression in female pups. These mitigating impacts were also evident in co-expression gene profiles that were enriched for estrogen-responsive genes. DNA methylation analyses indicated that maternal licking/grooming levels influenced the number of differentially methylated regions for prenatal bisphenol-exposed pups. These differentially methylated regions were enriched for binding sites for transcription factors that are known to interact with estrogen receptors. Finally, supplemental “licking-like” tactile stimulation normalized the DNA methylome in prenatal bisphenol-exposed female pups, with an increase and normalization of ESRRG binding to ESRRG-responsive genes.

**Conclusions:** These results identify a novel biological mechanism in which maternal licking/grooming can mitigate the negative neurodevelopmental impacts of prenatal bisphenol exposure and provides a proof-of-concept for postnatal tactile stimulation as a potential intervention strategy to improve neurodevelopmental outcomes.

## 1. Background

Early-life environmental exposures have a profound influence on neurodevelopmental trajectories and can lead to enduring changes in behavior. In naturalistic contexts, individuals are exposed to a combination of risk and protective factors that shape neurodevelopmental trajectories via distinct and common biological pathways. In particular, interactions between early-life environmental (e.g. pollutants, nutritional) and social experiences (e.g. deprivation or enhancement of maternal caregiving) appear to be prevalent in predicting risk and resilience to neurodevelopmental and mental health conditions [1,2]. The biological mechanisms underlying these interactions are largely unknown but are critical for developing effective intervention programs to mitigate the impacts of early-life adversity.

Prenatal exposure to endocrine disrupting chemicals has been shown in human and nonhuman animal studies to alter neurodevelopmental trajectories and increase the risk of neurodevelopmental and mental health conditions [3–5]. Bisphenols (BP) are commonly used plasticizers that are present in many plastics, such as food storage containers and reusable water bottles. Bisphenols such as BPA and other “BPA-free” structural analogs, including BPF and BPS, have also been shown to act as endocrine disruptors that primarily affect estrogen receptor signaling [6,7]. Previous studies have demonstrated that bisphenols can bind to estrogen receptor alpha (ESR1) and estrogen receptor beta (ESR2) to exert effects on gene transcription [6,8,9]. Prenatal exposure to bisphenols has been associated with negative neurodevelopmental and behavior outcomes for children in human epidemiological studies [10–16] as well as in rodent models of prenatal bisphenol exposure [17–22]. However, these prenatal BP effects may be attenuated by the experience of elevated levels of postnatal maternal care [23–26], suggesting the potential for opposing biological effects of these exposures. Identifying these biological pathways could facilitate the development of more targeted and effective interventions to mitigate the deleterious impacts of prenatal bisphenol exposure.

Prenatal bisphenol exposure [19,21,27,28] and variations in postnatal maternal care [29–32] are associated with changes in DNA methylation levels across the genome which can lead to stable changes in gene expression. There is accumulating evidence that steroid hormone receptor signaling links environmental exposures to DNA methylation modifications proximal to their respective transcription factor binding sites [33–37]. This includes estrogen receptor signaling, where estrogen receptor binding at estrogen responsive elements (EREs) has been associated with downstream changes in DNA methylation levels proximal to EREs [38–41]. Rats with perinatal BPA exposure have decreased transcript abundance of *Esr1* in the medial preoptic area (MPOA) of the hypothalamus later in life [42,43]; an effect observed particularly in females. Prenatal bisphenol exposure has also been associated with altered expression of estrogen-responsive genes with corresponding changes in DNA methylation levels within these genes [27,44]. Therefore, the effects of bisphenols on *Esr1* expression and estrogen receptor signaling at EREs may confer changes in DNA methylation levels at proximal CpG sites in the MPOA. Prenatal BPA exposure is also associated with changes in the expression of other estrogen and estrogen-related receptors, including *Esr2, Esrra, and Esrrg* [19,45–48], but it is currently unknown whether these changes occur in the MPOA specifically. Concurrently, high levels of postnatal maternal care received (licking/grooming or licking-like stimulation) has been associated with increased gene expression and decreased DNA methylation of *Esr1* throughout life in the MPOA [31,32,49]; an effect also occurring predominantly in females. Adult offspring with high levels of postnatal maternal licking/grooming received also have increased expression of *<u>Esr2</u>* and *Esrrg* in the hypothalamus [19]. Therefore, higher maternal licking/grooming received might mitigate the effects of prenatal bisphenol exposure on DNA methylation proximal to EREs later in life by normalizing estrogen and/or estrogen-related receptor expression levels and binding to DNA in the developing MPOA.

The primary objective of the current study was to examine the interactive effects of prenatal bisphenol exposure and postnatal maternal care on the transcriptome and DNA methylation modifications at specific transcription factor binding sites, notably EREs, in the developing MPOA. The MPOA may be especially vulnerable to prenatal bisphenol exposure because there is a high concentration of estrogen receptors in this brain region. In addition to profiling the MPOA transcriptome and methylome, we adapted a Suffix Array Kernel Smoothing (SArKS) correlative method to identify DNA motifs and potential transcription factor binding sites at differentially methylated regions. To determine the causal influence of maternal care on DNA methylation, we implemented a methodology involving postnatal “licking-like” tactile stimulation on DNA methylation modifications and estrogen receptor binding at proximal transcription factor binding sites. We hypothesized that postnatal maternal licking/grooming would mitigate the effects of prenatal bisphenol exposure on estrogen-responsive genes. Overall, we predicted that there would be sex-specific effects, with female offspring more sensitive to the interactive effects of prenatal bisphenol exposure and postnatal maternal licking/grooming on hypothalamic DNA methylation modifications.

## 2. Results

The experimental designs for the Maternal LG analyses are shown in **Figure 1A** and the Tactile LG analyses are shown in **Figure 1B**. The preprocessing and analysis pipelines for 3’ tag sequencing (tag-seq), oxidative reduced representation bisulfite sequencing (oxRRBS), and multi-omic factor analyses (MOFA) are summarized in **Figure 1C**. The full results, including the 50 μg/kg BPA and Mixed BP treatment groups in the Maternal LG analysis, can be found in the **Supplementary Results**.

**Figure 1.**
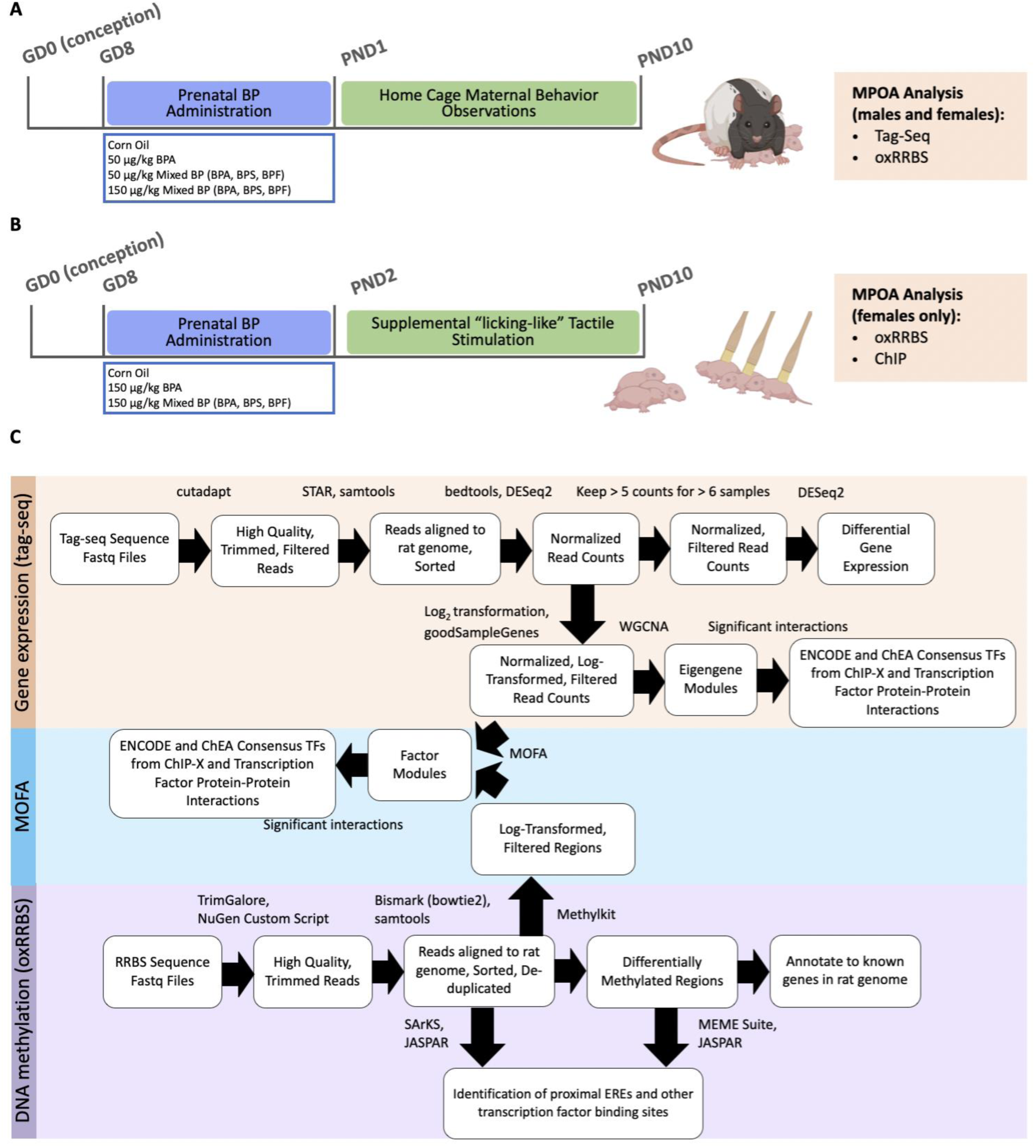
Study design and bioinformatic analysis pipelines. Experimental timeline for the Maternal LG analyses, which examined the effects of natural variations in maternal care (**A**). Experimental timeline for the Tactile LG analyses, which examined the effects of supplemental licking-like tactile stimulation (**B**). Preprocessing and analysis pipelines for gene expression, DNA methylation, and multi-omic analyses (**C**).

### 2.1 Limited effects of prenatal BP and postnatal maternal LG on differentially expressed genes (DEGs) in the MPOA

We performed genome-wide gene expression analyses using 3’ tag-sequencing to explore the effects of prenatal BP exposure and postnatal maternal care on estrogen receptor signaling. This analysis included examination of changes in the expression of estrogen receptor genes and estrogen-responsive genes. We first examined the main effects of prenatal BP and its interactions with postnatal maternal care on DEGs. **Supplementary Table S1** summarizes the DESeq2 output. Overall, there were very few DEGs found in all comparisons. The only exception was 63 DEGs (FDR ≤ 0.10) found when examining the interactions between the 150 μg/kg Mixed BP exposure group and maternal licking/grooming in female pups; however, none of these DEGs passed FDR correction of ≤ 0.05.

### 2.2 Interactive effects of prenatal BP and postnatal maternal LG on estrogen receptor expression in the MPOA

We next examined the main effects of prenatal BP and its interactions with postnatal maternal care on the expression levels of estrogen and estrogen-related receptors (*Esr1, Esr2, Esrra, Errb, Esrrg*), as there is previous work showing that both early-life environmental exposures impact estrogen receptor expression. We found a significant interaction between the 150 μg/kg Mixed BP exposure group and maternal licking/grooming on estrogen-related receptor gamma expression (*Esrrg*; p = 0.0004) in the female pups (**Figure 2A**) but not the male pups (**Figure 2B**). For female pups with 150 μg/kg Mixed BP exposure, maternal licking/grooming was negatively correlated with *Esrrg* expression while there was no strong relationship between maternal licking/grooming and *Esrrg* expression in the Corn Oil group. Female pups with 150 μg/kg Mixed BP exposure and high maternal licking/grooming had similar or lower *Esrrg* expression levels as the Corn Oil group. There were no other significant interactions with the other estrogen and estrogen-related receptors **(Supplementary Figure S2**). There were also no significant interactions found between prenatal bisphenol exposure and nest attendance.

**Figure 2.**
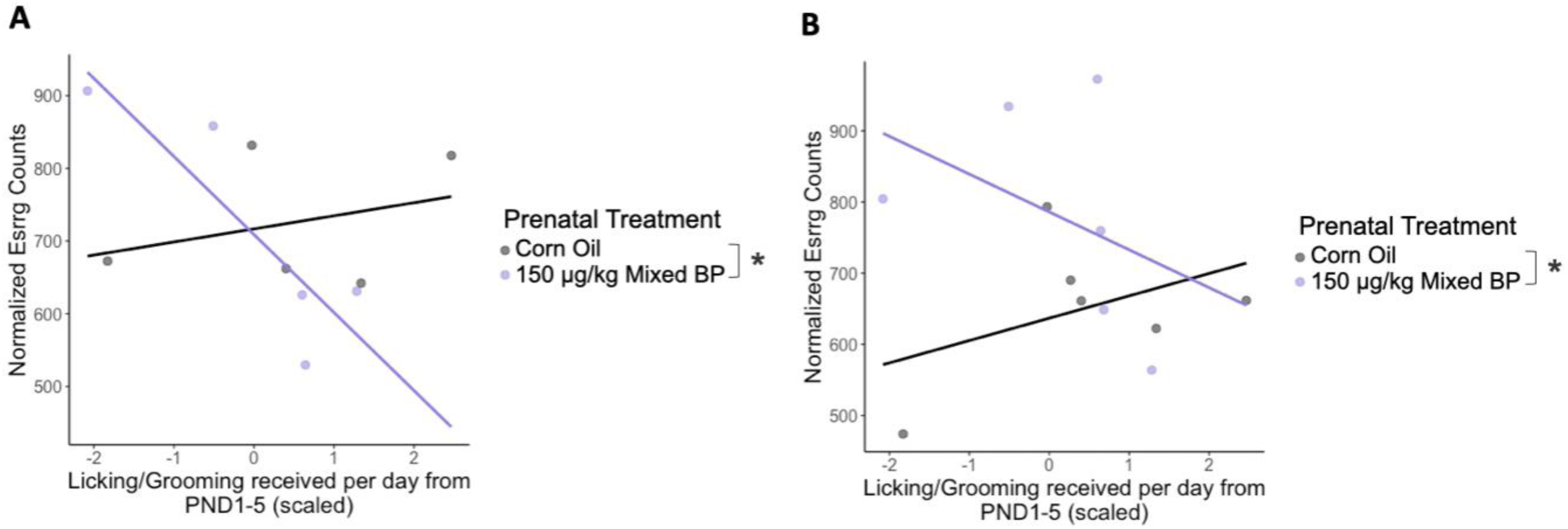
Sex-specific interaction between prenatal bisphenol exposure and postnatal maternal licking/grooming on *Esrrg* expression in the medial preoptic area. A significant interaction between the 150 μg/kg Mixed BP exposure group and maternal licking/grooming was found in female pups (**A**) but not male pups (**B**). In females from the 150 μg/kg Mixed BP group, maternal licking/grooming was negatively correlated with *Esrrg* expression. Scatterplots are displayed with linear regression lines for each prenatal treatment group. * p < 0.05 interaction between prenatal treatment and postnatal maternal care; # p < 0.10 interaction between prenatal treatment and postnatal maternal care

### 2.3 WGCNA modules associated with interactive effects of prenatal BP and postnatal maternal LG

The expression levels of individual genes are not independent from each other. Changes in gene expression tend to have a network-like structure of many co-expressed genes that are regulated by common transcription factors, such as estrogen receptors. Weighted Gene Co-expression Network Analysis (WGCNA) allows us to cluster these naturally occurring co-expressed gene networks into modules to examine the main effects of prenatal BP and its interactions with postnatal maternal care on differentially expressed gene networks. We can then examine if these gene modules are enriched in estrogen-responsive genes and their connectivity to estrogen and estrogen-related receptor expression. For the WGCNA analyses, a total of 7 modules for male pups and 13 modules for female pups were analyzed. For the female pups, there were significant or marginal interactions between the 150 μg/kg Mixed BP exposure group and maternal licking/grooming on five modules (Blue, Brown, Magenta, Turquoise, Yellow) with no significant main effects or interactions for male pups in the 150 μg/kg Mixed BP exposure group. More specifically, there were significant interactions for the Brown module (t = −2.544, p = 0.0292; 2,481 genes; **Figure 3A**), the Turquoise module (t = 2.291, p = 0.0449; 5,229 genes; **Figure 3B**), and the Yellow module (t = 2.061, p = 0.0264; 1,445 genes; **Figure 3C**). In females from the 150 μg/kg Mixed BP group, maternal licking/grooming was negatively correlated with the Brown eigengene values and positively correlated with the Turquoise and Yellow eigengene values. In the Corn Oil group, the relationships between maternal licking/grooming and the WGCNA module eigengene values were reversed. There were also marginal interactions for the Blue module (t = −2.179, p = 0.0544; 4,022 genes; **Supplemental Figure S1A**) and the Magenta module (t = −2.205, p = 0.0520; 980 genes; **Supplemental Figure S1B**), Top GO terms for each of the modules with significant interactions in female offspring are presented in **Figure 3D**; the top GO terms for the modules with marginal interactions are presented in **Supplemental Figure S1C**. There were no significant interactions found between prenatal bisphenol exposure and nest attendance; however, including nest attendance and the interaction between prenatal bisphenol exposure and nest attendance in the linear model was required to reveal the interactions between the 150 μg/kg Mixed BP exposure group and maternal licking/grooming.

**Figure 3.**
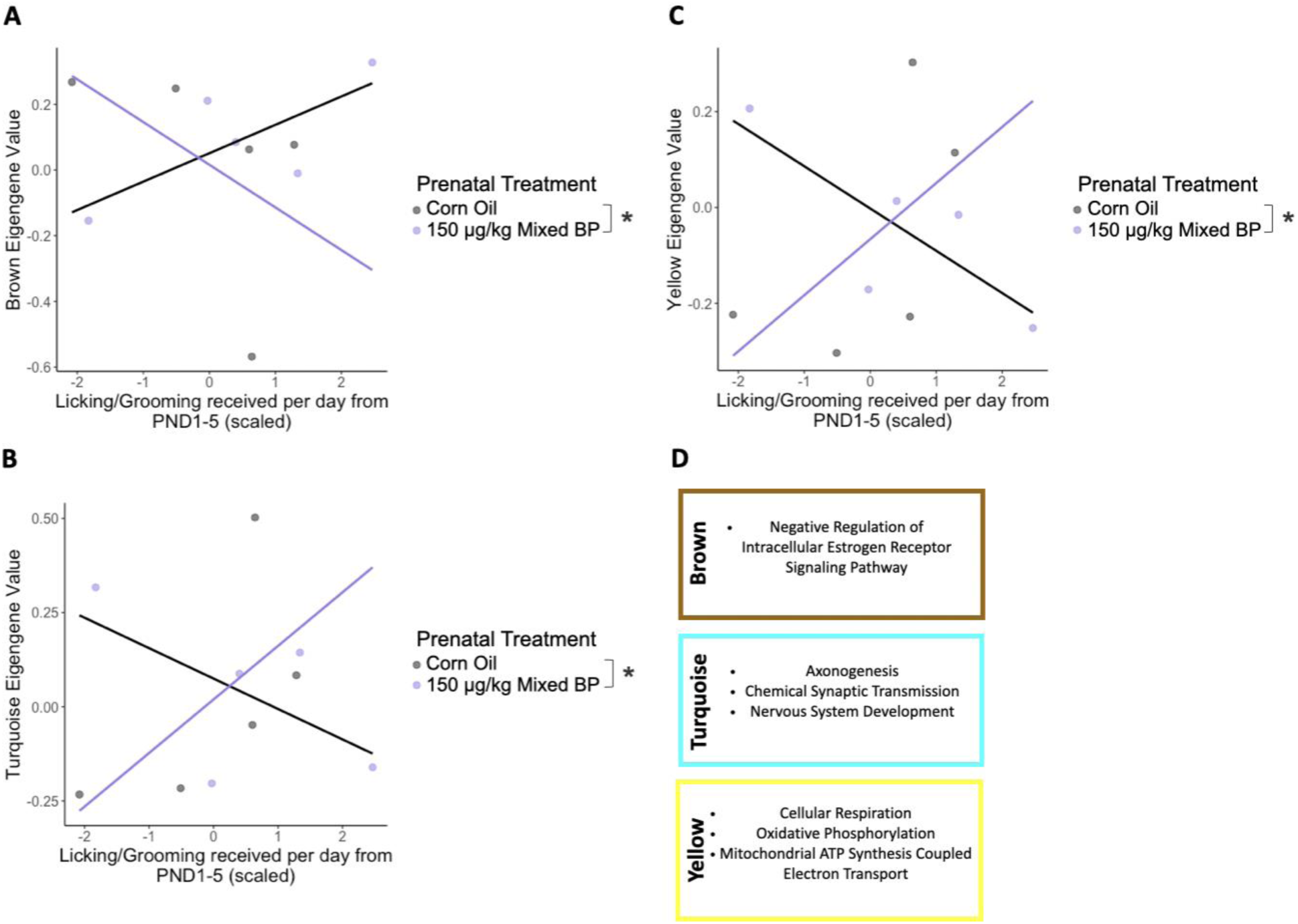
Interactions between prenatal bisphenol exposure and postnatal maternal licking/grooming on co-expressed gene modules in the medial preoptic area of female pups. Significant interactions between the 150 μg/kg Mixed BP exposure group and maternal licking/grooming were found in the Brown (**A**), Turquoise (**B**), and Yellow (**C**) eigengene modules. In females from the 150 μg/kg Mixed BP group, maternal licking/grooming was negatively correlated with the Brown eigengene values and positively correlated with the Turquoise and Yellow eigengene values. Top GO terms (**D**) from these modules were related to estrogen receptor signaling, neurodevelopment, neurotransmission, and cellular metabolism. Scatterplots are displayed with linear regression lines for each prenatal treatment group. * p < 0.05 interaction between prenatal treatment and postnatal maternal care

### 2.4 Gene expression module enrichment for estrogen-responsive genes and genes that interact with estrogen receptors

Among the modules in which there were significant interactions between prenatal bisphenol exposure and postnatal maternal care in female offspring (**Figure 3**), we examined enrichment for estrogen-responsive genes in the ENCODE and ChEA Consensus TFs from ChIP-X module using Enrichr. We also examined enrichment for genes that interact with estrogen receptors since our prior work using WGCNA in developing prefrontal cortex and amygdala has found strong enrichment of ESR1 in the Transcription Factor Protein-Protein Interactions module. In the Brown module, there was no enrichment of ESR1 found in the list of ENCODE and ChEA Consensus TFs from ChIP-X or Transcription Factor Protein-Protein Interactions. Notably, this module contains *Esr1*, *Esr2*, and *Esrrg*. In the Turquoise module, there was a marginal enrichment of ESR1 found in the list of ENCODE and ChEA Consensus TFs from ChIP-X (FDR = 0.054) and a significant enrichment of ESR1 found in the list of Transcription Factor Protein-Protein Interactions (FDR = 0.003). In the Yellow module, there was a significant enrichment of ESR1 found in the list of ENCODE and ChEA Consensus TFs from ChIP-X (FDR = 0.038) and Transcription Factor Protein-Protein Interactions (FDR < 0.001).

### 2.5 Connectivity between estrogen receptors and all gene expression modules with interactions between prenatal BP and postnatal maternal LG

To further connect the differentially co-expressed gene modules to estrogen receptor signaling changes, we examined the connectivity of each of the estrogen and estrogen-related receptors for each module in which there were significant interactions between prenatal BP and postnatal maternal care in female offspring. **Table 1** summarizes these results. Most notably, *Esrrg* had moderate-high positive connectivity with the Brown module (73%) and moderate-high negative connectivity with the Turquoise and Yellow modules (−69% and −54%, respectively).

**Table 1.** Connectivity (kME) of estrogen receptors with genes within the co-expressed gene modules generated by the WGCNA analysis. Values are represented as percentages of genes within each module that have significant correlations. Negative values indicate inverse correlations.

|  | kME Brown | kME Turquoise | kME Yellow |
| --- | --- | --- | --- |
| *Esr1* | 32.93 | -13.40 | -1.53 |
| *Esr2* | 41.06 | -34.39 | -48.27 |
| *Esrra* | 18.76 | -20.91 | -14.89 |
| *Esrrb* | 25.80 | -54.04 | -17.94 |
| *Esrrg* | 73.13 | -68.89 | -53.58 |

### 2.6 Impact of postnatal maternal LG on differentially methylated regions in prenatal BP-exposed pups

Given that there were strong indications that the effects of prenatal BP exposure and postnatal licking/grooming converge on estrogen receptor signaling, we next examined the influence of postnatal maternal licking/grooming on differentially methylated regions (DMRs) in the developing MPOA among the prenatal treatment groups. To do so, we assessed the degree of overlap in DMRs when we include or exclude maternal licking/grooming as a covariate while comparing DNA methylation levels between the 150 μg/kg Mixed BP exposure group to the Corn Oil group. Overall, while many DMRs were unchanged when including maternal licking/grooming as a covariate, there were some discrepancies found for both male and female pups. Notably, there were more DMRs that were added than lost when including maternal licking/grooming as a covariate.

For males, there were 44 DMRs (corresponding to 27 genes) that were unchanged, 6 DMRs (corresponding to four genes) that were lost, and 38 DMRs (corresponding to 21 genes) that were added when including maternal licking/grooming as a covariate (**Figure 4A**; **Supplementary Table S4**). For females, there were 60 DMRs (corresponding to 48 genes) that were unchanged, 10 DMRs (corresponding to eight genes) that were lost, and 25 DMRs (corresponding to 16 genes) that were added when including maternal licking/grooming as a covariate (**Figure 4B; Supplementary Table S7**).

**Figure 4.**
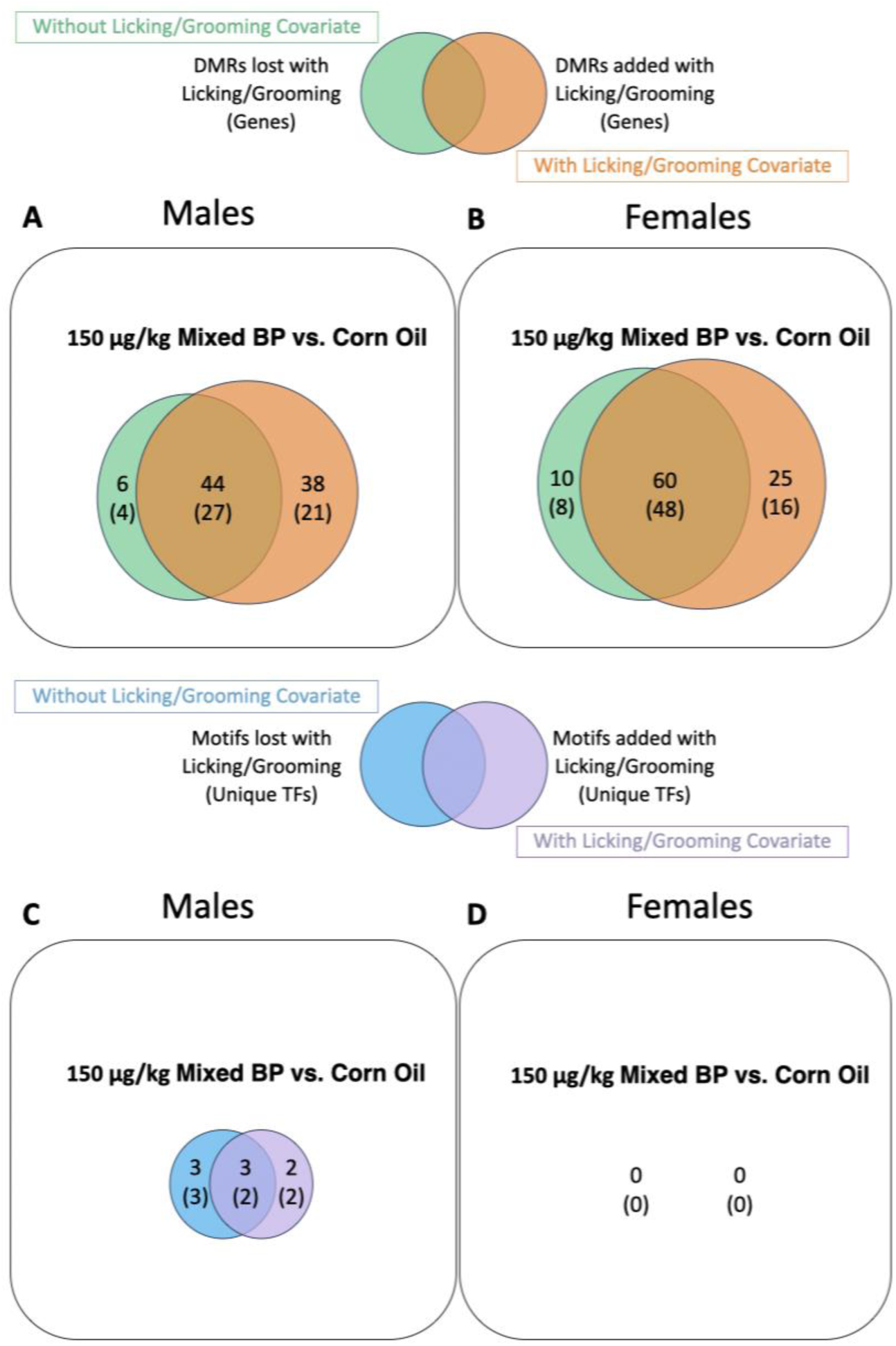
Influence of postnatal maternal licking/grooming on differentially methylated regions (DMRs) and differentially methylated transcription factor binding sites of male and female pups with prenatal 150 μg/kg Mixed BP exposure. Including maternal licking/grooming as a covariate altered DMRs in both male (**A**) and female (**D**) pups. For both comparisons, more DMRs were added than lost when adding licking/grooming as a covariate. Using the SArKS method, including maternal licking/grooming as a covariate altered overrepresented sequence motifs and predicted transcription factor binding sites in male pups (**C**) but no motifs or predicted transcription factor binding sites were identified in female pups (**D**). Venn diagrams (**A-B**) are displayed with the number of DMRs and annotated genes when excluding (green) and including (orange) maternal licking/grooming as a covariate. DMRs that do not overlap are considered to be influenced by postnatal maternal licking/grooming. Venn diagrams (**C-D**) are displayed with the number of motifs and unique transcription factors when excluding (blue) and including (purple) maternal licking/grooming as a covariate. Motifs that do not overlap are considered to be influenced by postnatal maternal licking/grooming.

### 2.7 Impact of postnatal maternal LG on differentially methylated transcription factor binding sites in prenatal BP-exposed pups

To connect the changes in estrogen receptor signaling found in the tag-seq data and DNA methylation changes more directly, we examined the influence of postnatal maternal licking/grooming on differential methylation of proximal transcription factor binding sites and EREs. To assess this outcome, we performed a motif analysis using two different methods (SArKS and MEME Suite) to identify overrepresented sequence motifs and JASPAR to match the motifs to known transcription factor binding sites.

When <u>excluding</u> licking/grooming as a covariate in male pups with prenatal 150 μg/kg Mixed BP exposure, SArKS identified six significant motifs associated with five unique transcription factors in hypomethylated regions (**Supplementary Table S8**). When <u>including</u> licking/grooming as a covariate, SArKS identified four significant motifs associated with four unique transcription factors in hypomethylated regions (**Supplementary Table S9**). Overall, when including maternal licking/grooming as a covariate, two new transcription factor binding sites (EGR3, E2F7) were identified in hypomethylated regions (**Figure 4C**). Surprisingly, SArKS did not identify any significant motifs for female pups with prenatal 150 μg/kg Mixed BP exposure, either including or excluding maternal licking/grooming as a covariate (**Figure 4D**).

There were also changes in motifs and predicted transcription factor binding sites when adding postnatal maternal licking/grooming as a covariate with MEME Suite (**Supplementary Figure S2**). In many comparisons, there were a larger number of motifs and predicted transcription factor binding sites that were altered with postnatal maternal licking/grooming and less overlap using the MEME Suite method than the SArKS method. There were also several motifs and predicted transcription factor binding sites identified when including and excluding maternal licking/grooming as a covariate for female pups with prenatal 150 μg/kg Mixed BP exposure using the MEME Suite method (**Supplementary Figure S2).**

While no estrogen receptors were found in the list of predicted transcription factor binding sites, a portion of the predicted transcription factors found in male pups have known interactions with estrogen receptors. This includes transcription factors that are known to dimerize with ESR1, block estrogen receptors from binding to DNA, facilitate transcription of *Esr1*, or are estrogen-responsive genes themselves (**Table 2**).

**Table 2.** List of unique transcription factors associated with prenatal bisphenol exposure and/or postnatal maternal licking/grooming in male pups and their known interactions with estrogen receptors.

| Analysis | Methylation comparison | Transcription factor name | Relationship to estrogen receptors |
| --- | --- | --- | --- |
| **Unchanged with Licking/Grooming** |  |  |  |
| 150 µg/kg Mixed BP vs. Corn Oil | Hypomethylated | ZNF417 |  |
| 150 µg/kg Mixed BP vs. Corn Oil | Hypomethylated | SP5 |  |
| **Lost with Licking/Grooming** |  |  |  |
| 150 µg/kg Mixed BP vs. Corn Oil | Hypomethylated | EBF2 |  |
| 150 µg/kg Mixed BP vs. Corn Oil | Hypomethylated | **HAND2** | [50,51] |
| 150 µg/kg Mixed BP vs. Corn Oil | Hypomethylated | **RBPJ** | [52] |
| **Added with Licking/Grooming** |  |  |  |
| 150 µg/kg Mixed BP vs. Corn Oil | Hypomethylated | **EGR3** | [53] |
| 150 µg/kg Mixed BP vs. Corn Oil | Hypomethylated | E2F7 |  |

### 2.8 Interactive effects of prenatal BP exposure and postnatal maternal LG on the multi-omic epigenome in the MPOA of female pups

The disconnect between gene expression and DNA methylation changes may be in part due to the different analysis methods used to examine the interactions between prenatal BP exposure and maternal care. To address this disparity, we also examined the influence of postnatal maternal licking/grooming on the combined DNA methylome and transcriptome in the developing MPOA using Multi-Omic Factor Analysis (MOFA). For female pups, there was one factor (Factor 4) that showed significant interactions between the 150 μg/kg Mixed BP exposure group and maternal licking/grooming (t = 4.456, p = 0.0043; **Figure 5A**) as well as nest attendance (t = 3.182, p = 0.0190; **Figure 5B**). Factor 4 values were more strongly positively correlated with maternal licking/grooming in females from the 150 μg/kg Mixed BP group than the Corn Oil group. Factor 4 values were negatively correlated with nest attendance in the 150 μg/kg Mixed BP group, though the smaller range of nest attendance values limit full interpretation of the data. In the Corn Oil group, Factor 4 values were positively correlated with nest attendance. Similar to the WGCNA analysis, including nest attendance and the interaction between prenatal bisphenol exposure and nest attendance in the linear model was required to reveal the significant interaction between the 150 μg/kg Mixed BP exposure group and maternal licking/grooming.

**Figure 5.**
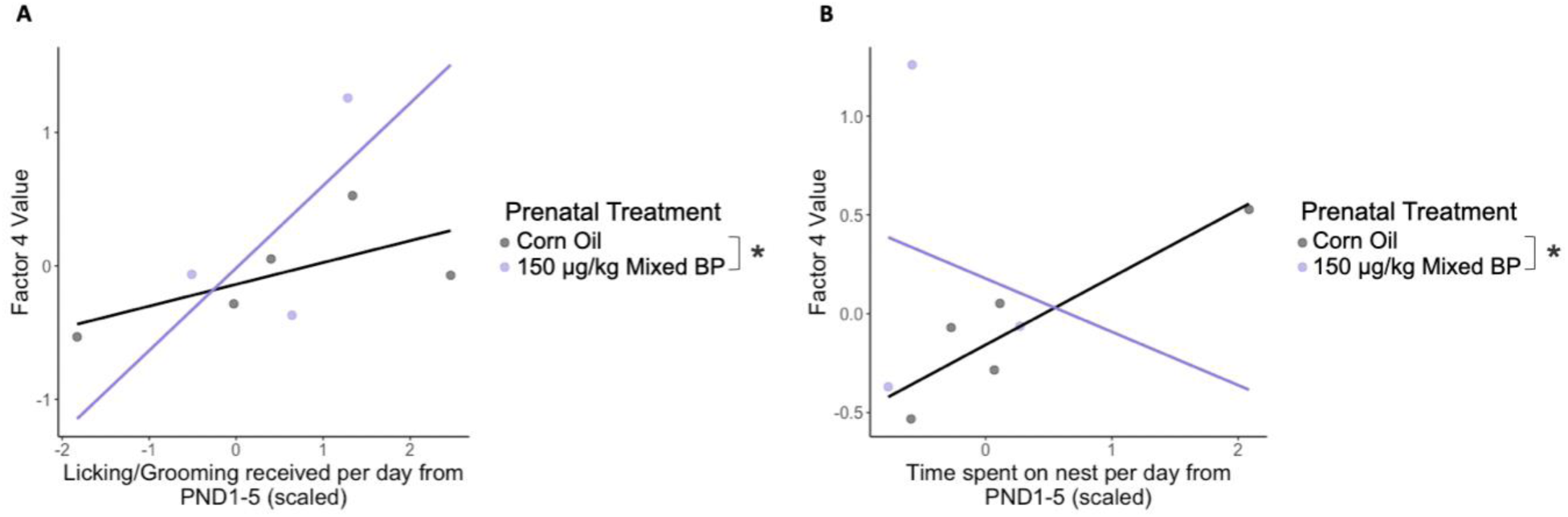
Interactions between prenatal bisphenol exposure and postnatal maternal licking/grooming on Factor 4 generated from the multi-omic factor analysis (MOFA) in female pups. Significant interactions between the 150 μg/kg Mixed BP exposure group and maternal licking/grooming (**A**) and nest attendance (**B**) were found when combining the transcriptome and DNA methylome datasets. In females from the 150 μg/kg Mixed BP group, Factor 4 values were positively correlated with maternal licking/grooming and negatively correlated with nest attendance. Scatterplots are displayed with linear regression lines for each prenatal treatment group. * p < 0.05 interaction between prenatal treatment and postnatal maternal care

#### 2.8.1 Factor module enrichment for estrogen-responsive genes and genes that interact with estrogen receptors

For Factor 4, there were only 14 genes that overlapped between the top genes in the tag-seq and oxRRBS datasets. For the top genes in the tag-seq dataset (467 genes analyzed) there was a significant enrichment of ESR1 found in the list of Transcription Factor Protein-Protein Interactions (FDR < 0.001). For the top genes in the oxRRBS dataset (423 genes analyzed) there was a significant enrichment of ESR1 found in the list of ENCODE and ChEA Consensus TFs from ChIP-X (FDR = 0.031). There was also a nominal enrichment of ESR1 found in the list of Transcription Factor Protein-Protein Interactions (p = 0.0178), but it did not pass FDR correction (FDR > 0.10).

### 2.9 Impact of supplemental tactile stimulation on differentially methylated regions in prenatal bisphenol-exposed female pups

The findings so far indicated that higher postnatal licking/grooming can mitigate the impacts of prenatal bisphenol exposure on ESRRG signaling in female pups. However, the corresponding changes in DNA methylation were less clear. Postnatal licking/grooming can co-occur with other maternal behaviors such as thermotactile contact, which can also impact DNA methylation [37], so establishing the causality of licking-like tactile stimulation in mitigating the impacts of prenatal bisphenol exposure on ESRRG signaling and DNA methylation changes is important. Therefore, we examined the impacts of experimentally induced increases in licking-like tactile stimulation on the MPOA DNA methylome in prenatal bisphenol-exposed female pups. We also examined the influence of the bisphenol mixture used by including a BPA-only exposure group in the subsequent analyses. For nonstimulated pups, there were 113 DMRs (63 hypermethylated, 50 hypomethylated; **Supplementary Table S13**) between the 150 μg/kg BPA exposure group and Corn Oil group while there were 246 DMRs (227 hypermethylated, 19 hypomethylated; **Supplementary Table S14**) between the 150 μg/kg Mixed BP exposure group and Corn Oil group. There were notable reductions in DMRs for the stimulated siblings—46 DMRs (34 hypermethylated, 12 hypomethylated; **Supplementary Table S15**) between the 150 μg/kg BPA exposure group and Corn Oil group and 22 DMRs (12 hypermethylated, 10 hypomethylated; **Supplementary Table S16**) between the 150 μg/kg Mixed BP exposure group and Corn Oil group (**Figure 6A**). This equates to about a 60% reduction and over 90% reduction in DMRs in the 150 μg/kg BPA and 150 μg/kg Mixed BP exposure groups, respectively. There were only 27 DMRs between nonstimulated and stimulated siblings in the Corn Oil group, so the reductions in DMRs in the prenatal bisphenol treatment groups are unlikely due to differences between nonstimulated and stimulated pups in the Corn Oil group per se.

**Figure 6.**
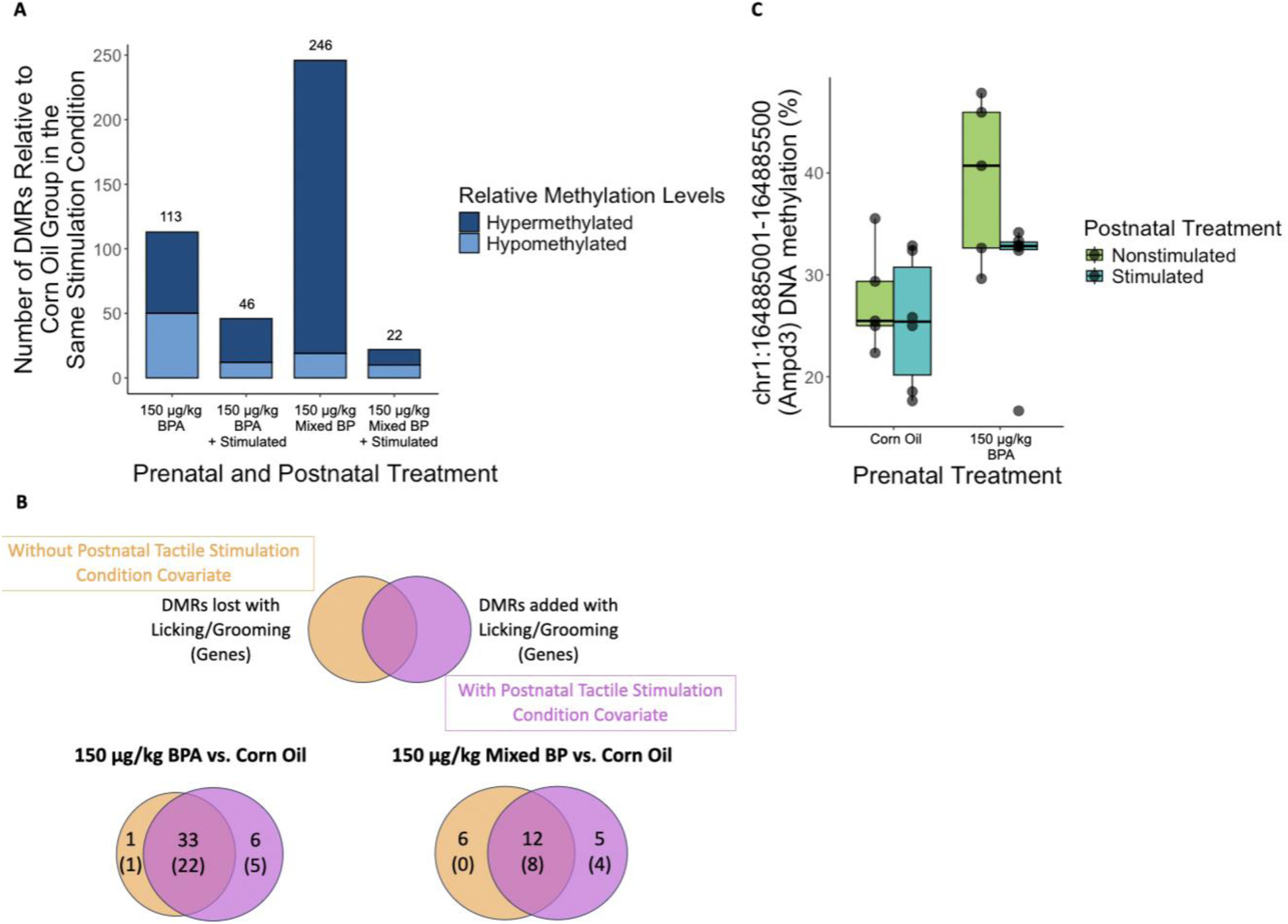
Supplemental tactile stimulation normalizes the DNA methylome genome-wide in the prenatal bisphenol-exposed female pup MPOA. Female pups with supplemental tactile stimulation showed a marked reduction (60-90%) in DMRs relative to their nonstimulated siblings when comparing prenatal bisphenol-exposed pups to the Corn Oil group in their respective tactile stimulation condition (**A**). Hypermethylation in the 150 μg/kg Mixed BP exposure group appears to be particularly affected. However, we found much fewer DMRs when reanalyzing the dataset by comparing the prenatal bisphenol exposure group and Corn Oil group with and without including postnatal tactile stimulation condition as a covariate (**B**). One DMR, annotated to the promoter region of the Ampd3 gene, showed a significant interaction between the 150 μg/kg BPA exposure group and postnatal tactile stimulation condition via a mitigating effect of supplemental tactile stimulation in the 150 μg/kg BPA exposure group (**C**). Boxplot with the median +/-interquartile range are displayed with individual datapoints.

However, a more complex picture emerges when examining the effects of supplemental tactile stimulation within the prenatal bisphenol treatment groups and the interactions between prenatal treatment group and postnatal tactile stimulation condition. We only found 14 DMRs (8 hypermethylated and 6 hypomethylated) between the nonstimulated and stimulated siblings in the 150 μg/kg BPA exposure group and 32 DMRs (3 hypermethylated and 29 hypomethylated) between the nonstimulated and stimulated siblings in the 150 μg/kg Mixed BP exposure group. When combining the nonstimulated and stimulated siblings into one analysis, there were 34 DMRs between the 150 μg/kg BPA exposure group and Corn Oil group and 18 DMRs between the 150 μg/kg Mixed BP exposure group and Corn Oil group. Adding postnatal tactile stimulation condition as a covariate revealed six additional DMRs and one lost DMR in the 150 μg/kg BPA exposure group and five additional DMRs and six DMRs lost in the 150 μg/kg Mixed BP exposure group (**Figure 6B**). Upon further inspection of the DMRs that were added when including postnatal tactile stimulation condition as a covariate, there was only one DMR where the nature of the interaction was a mitigating effect of supplemental tactile stimulation in the 150 μg/kg BPA exposure group (**Figure 6C**). This DMR was annotated to the promoter region of Adenosine Monophosphate Deaminase 3 (Ampd3), which is a known estrogen-responsive gene [54]. Overall, while we found widespread reductions in DMRs across the genome with supplemental tactile stimulation, the effect sizes for the individual DMRs were small. In addition, we observed a bias toward hypermethylation in the 150 μg/kg Mixed BP exposure group that was normalized with supplemental tactile stimulation.

### 2.10 Impact of supplemental tactile stimulation on ESRRG binding to candidate genes in prenatal bisphenol-exposed female pups

Given that supplemental tactile stimulation can mitigate the effects of prenatal BP exposure on the DNA methylome in female pups, we next examined the role of ESRRG signaling changes in a more direct way by measuring ESRRG binding to DNA. In our DMR lists, we identified four known ESRRG-responsive genes based on a previous ChIP-seq study with mouse neurons [55]. This includes Enolase 4 (Eno4; **Figure 7A**), Forkhead Box P1 (Foxp1; **Figure 7B**), Nuclear Receptor Corepressor 2 (Ncor2; **Figure 7C**), and Protein Phosphatase 1 Regulatory Subunit 3B (Ppp1r3b; **Figure 7D**). We focused on these known ESRRG-responsive genes that were hypermethylated in nonstimulated pups in the 150 μg/kg BPA exposure group (Ppp1r3b; **Figure 7D**) and 150 μg/kg Mixed BP exposure group (Eno4; **Figure 7A**, Foxp1; **Figure 7B**) but were not differentially methylated in the stimulated pups. As a negative control, we also examined one ESRRG-responsive gene that was hypermethylated in stimulated pups in the 150 μg/kg BPA exposure group (Ncor2; **Figure 7C**). Finally, we examined one putative ERE in intron 5 of the Septin9 gene since this DMR was hypermethylated in nonstimulated pups in both the 150 μg/kg BPA and 150 μg/kg Mixed BP exposure groups (and not differentially methylated in the stimulated pups; **Figure 7E**) and Septin9 appears in one of the DMR lists of our previous oxRRBS dataset examining the effects of natural variations in licking/grooming (**Supplementary Table S2**).

**Figure 7.**
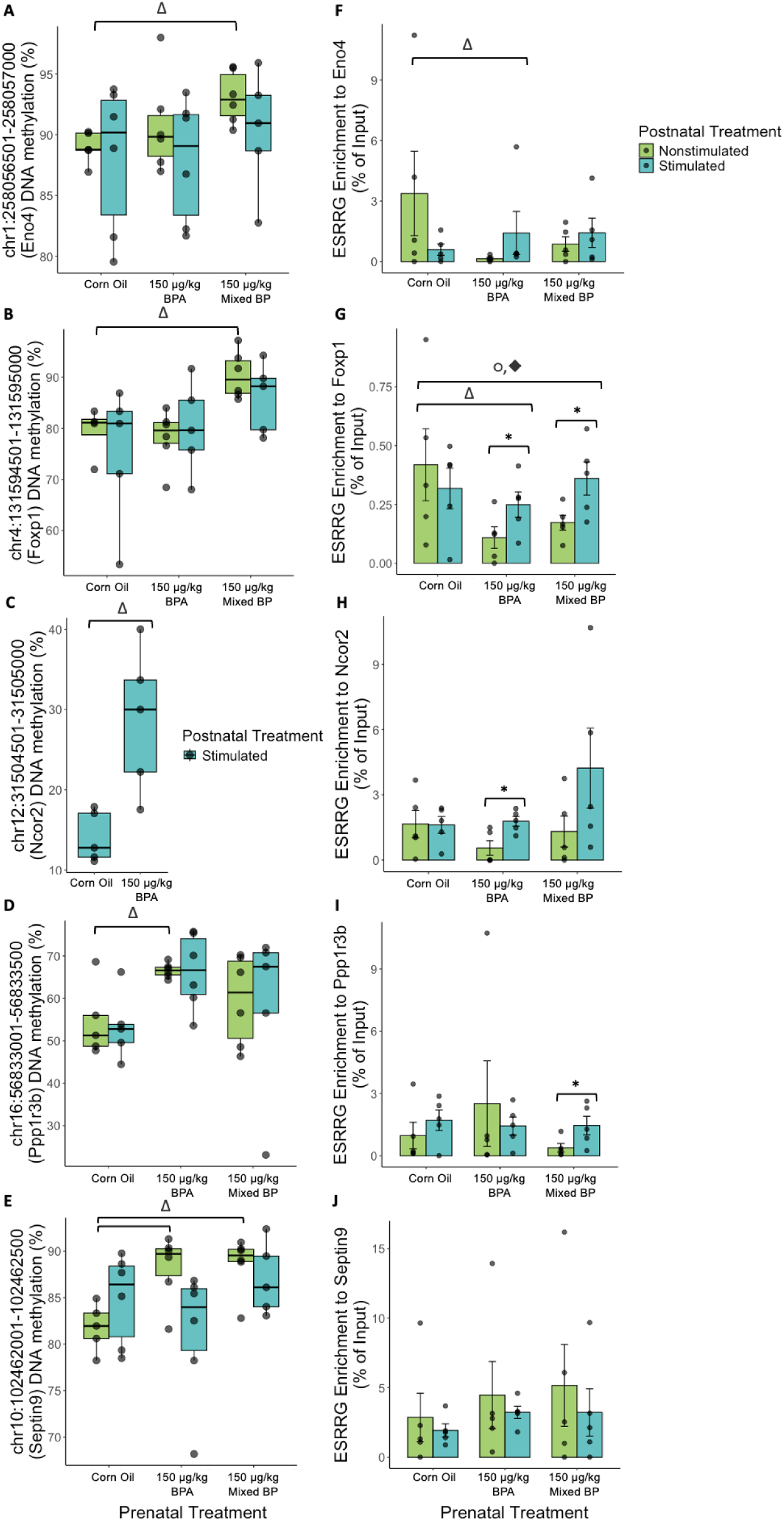
Supplemental tactile stimulation normalizes ESRRG binding to ESRRG-responsive genes in the prenatal bisphenol-exposed female pup MPOA. Several ESRRG-responsive genes were identified in the oxRRBS DMR datasets, including Eno4 (**A**), Foxp1 (**B**), Ncor2 (**C**), and Ppp1r3b (**D**). Only the differences in methylation between the corn oil and 150 μg/kg BPA exposure group in stimulated pups for Ncor2 (**C**) are displayed because there was insufficient coverage to analyze the other treatment groups. Septin9 (**E**) showed hypermethylation in both the 150 μg/kg BPA and 150 μg/kg Mixed BP exposure groups in the nonstimulated pups and contained a classical ERE sequence in intron 5. Chromatin immunoprecipitation of ESRRG revealed a reduction in ESRRG binding in prenatal bisphenol-exposed pups that was normalized in the stimulated pups for the ESRRG-responsive genes (**F, G, H, I**). A significant interaction between the 150 μg/kg Mixed BP exposure group and postnatal tactile stimulation condition was found for Foxp1 specifically (**G**). Septin9 showed a different pattern of ESRRG binding (**J**) with no significant differences or interactions between prenatal or postnatal treatment groups. Boxplots (**A, B, C, D, E**) with the median +/- interquartile range and bar graphs (**F, G, H, I, J**) with mean +/- standard error are displayed with individual datapoints. Δ p or q < 0.05 main effect of prenatal treatment; ○ p < 0.10 main effect of prenatal treatment; * p < 0.05 main effect of postnatal tactile stimulation condition within prenatal treatment group; ◆ p < 0.05 interaction between prenatal treatment group and postnatal tactile stimulation condition

Overall, we found that prenatal bisphenol-exposed female pups had reduced ESRRG binding at the DNA binding motif of ESRRG-responsive genes, which was normalized for female pups with supplemental tactile stimulation. There was a significant main effect of prenatal treatment group for Eno4 and Foxp1, where there was a significant reduction of ESRRG binding between the Corn Oil group and 150 μg/kg BPA exposure group for Eno4 (t = −2.15, p = 0.044; **Figure 7F**) and Foxp1 (t = −2.63, p = 0.015; **Figure 7G**) and a marginal reduction between the Corn Oil group and 150 μg/kg Mixed BP exposure group for Foxp1 (t = −1.91, p = 0.07; **Figure 7G**). There were significant main effects of tactile stimulation within the 150 μg/kg BPA exposure group for Foxp1 and Ncor2, where the stimulated pups had significantly higher ESRRG binding for Foxp1 (t = 7.08, p < 0.001; **Figure 7G**) and Ncor2 (t = 3.13, p = 0.021; **Figure 7H**). There were also significant main effects of tactile stimulation within the 150 μg/kg Mixed BP exposure group for Foxp1 and Ppp1r3b, where the stimulated pups had significantly higher ESRRG binding for Foxp1 (t = 2.42, p = 0.042; **Figure 7G**) and Ppp1r3b (t = 2.83, p = 0.047; **Figure 7I**). There were no main effects of supplemental tactile stimulation for any of the genes tested in the Corn Oil group. Finally, there was a significant interaction between prenatal treatment and postnatal tactile stimulation condition for Foxp1 in the 150 μg/kg Mixed BP exposure group (t = 2.09, p = 0.049; **Figure 7G**). There were no main effects of prenatal treatment or postnatal tactile stimulation condition or any interactions for ESRRG binding for the Septin9 gene, though overall enrichment at the ERE was relatively high (**Figure 7J**).

## 3. Discussion

In this study, we examined the interacting and possibly mitigating impacts of postnatal maternal licking/grooming in prenatal bisphenol-exposed pups on the transcriptome and the DNA methylome in the developing MPOA. We also examined the mitigating impact of experimentally induced increases in licking-like tactile stimulation on the DNA methylome in the developing MPOA. Finally, we investigated the role of estrogen receptor signaling changes as a potential biological mechanism underlying these interactions. Overall, we found that postnatal maternal licking/grooming has a substantial influence on estrogen receptor signaling and DNA methylation changes, particularly in female pups. This might be driven by the mitigating effect of maternal licking/grooming on *Esrrg* expression and ESRRG binding activity throughout the genome. Given that ESRRG has a high affinity for bisphenols [56,57] and can facilitate gene expression by binding to estrogen responsive elements or its own specific binding motif, these results suggest a novel biological mechanism in which postnatal maternal care can mitigate the negative neurodevelopmental impacts of prenatal bisphenol exposure. Moreover, these results indicate that postnatal supplemental tactile stimulation may be an effective intervention against the adverse neurodevelopmental consequences of prenatal bisphenol exposure, particularly in females.

We hypothesized that the interactions between prenatal bisphenol exposure and postnatal maternal licking/grooming would primarily involve changes in *Esr1* expression in the MPOA and that the effects would be stronger in female pups than male pups. However, we found sex-specific interactions in *Esrrg* expression that appear to be associated with multiple co-expressed gene modules in the WGCNA analysis. While it is known that BPA exposure can stimulate expression of *Esrrg* in a sex-specific manner [19,48] and has a high binding affinity to ESRRG [56,57], less is known about the effects of postnatal maternal care on *Esrrg* expression or signaling changes. A prior study examining prenatal BPA exposure and postnatal maternal care conducted in our lab found brain region and sex-specific effects of maternal care on *Esrrg* expression, where there was an increase in expression in the prefrontal cortex in female offspring only and in the hypothalamus in male and female offspring [19]. There were no interactions between prenatal BPA and postnatal maternal care on *Esrrg* expression noted in the study. In our current study, we found postnatal maternal licking/grooming decreased and normalized *Esrrg* expression in the female pups with prenatal 150 μg/kg Mixed BP exposure. The discrepancies may be primarily due to the different ages assessed (weanlings vs. PND10 pups), the bisphenol doses used, and the bisphenol mixture used in our study appears to have a different effect than BPA alone.

While the sex-specific interactions in gene expression were consistent in our hypothesis, we also found that postnatal maternal licking/grooming has a substantial influence on DNA methylation changes in prenatal bisphenol-exposed male offspring. We also found in prior studies that postnatal maternal care can interact and mitigate the impacts of prenatal bisphenol exposure on later-life phenotypes and gene expression in the prefrontal cortex and amygdala at PND10 in male offspring [26]. Therefore, it is possible that postnatal maternal licking/grooming can mitigate the impacts of prenatal bisphenol exposure in male offspring, but the underlying biological mechanisms are currently unclear and/or are not strongly related to estrogen receptor signaling. As an example, previous work using another estrogenic endocrine disrupting chemical (Aroclor 1221) found estradiol levels predicted behavioral phenotypes in prenatal Aroclor 1221-exposed females while dopamine-related gene expression predicted behavioral phenotypes in prenatal Aroclor 1221-exposed male offspring [58]. These sex-specific impacts are worth exploring in future studies.

Our WGCNA analysis indicated that including postnatal nest attendance in our statistical models was required to unveil the significant interactions between the 150 μg/kg Mixed BP exposure group and maternal licking/grooming but was not significantly associated with these modules by itself. Postnatal nest attendance includes licking/grooming and other forms of pup contact, including thermotactile contact. This suggests that postnatal licking/grooming is the primary driver of these interactions and overall pup contact plays a minor but possibly important role [37] in gene expression changes for prenatal bisphenol-exposed female pups. This was confirmed in our subsequent experiments, which directly manipulated levels of licking-like tactile stimulation while keeping all pups in thermoneutral conditions.

Our initial DMR analyses indicate postnatal maternal licking/grooming has a substantial influence on DNA methylation changes in prenatal bisphenol-exposed pups. The finding that more DMRs were added than lost when including postnatal maternal licking/grooming as a covariate suggests that postnatal maternal licking/grooming plays more of a moderating role in shaping the DNA methylome in prenatal bisphenol-exposed pups, consistent with our prior findings [26,59]. However, it was apparent that the differential methylation findings did not map consistently onto differential gene expression. While it is assumed that changes in DNA methylation and gene expression would be strongly linked, there are several reasons why there may be a discordance. First, we did not implement any substantial environmental challenges to the pups immediately prior to brain collection.

While DNA methylation changes are relatively stable, gene expression changes are dependent on both the availability of chromatin to allow transcription and transcriptional activity itself. Therefore, the associations between DNA methylation and gene expression changes may be stimulus-dependent. The second reason is that there are inherent differences in the approach in which these changes are assessed. While we assessed the whole transcriptome, we only probed a small portion of CpG sites in the genome. In addition, many of these CpG sites are intergenic and we have an incomplete understanding of the role of DNA methylation in distal CpG sites on gene expression. Finally, in any particular cell, DNA methylation is mainly binary (methylated or unmethylated with rare instances of hemi-methylation) while gene expression changes can have a large range. This could complicate analyses when using bulk tissue and would need to be resolved at the single-cell level. Nevertheless, when we integrated the DNA methylation and gene expression data using MOFA, we generally replicate the gene expression findings. This confirms that the significant interactions between the 150 μg/kg Mixed BP exposure group and maternal licking/grooming in female pups apply to both gene expression and DNA methylation.

We explored potential transcription factor binding sites associated with DNA methylation changes with prenatal bisphenol exposure and postnatal maternal licking/grooming. To do so, we adapted the SArKS method to RRBS data so we can correlate DNA methylation changes in prenatal bisphenol-exposed pups with and without postnatal maternal licking/grooming as a covariate to motifs and possible transcription factor binding sites. As a comparison, we also used MEME Suite to identify motifs based on the changes in DMRs with and without postnatal maternal licking/grooming as a covariate. The main difference with using a correlative-based method is that it accounts for the continuous range of DNA methylation changes (0-100%) instead of a binary representation (i.e., it is differentially methylated or not differentially methylated). A correlative-based method can be particularly advantageous with more complex study designs where multiple independent variables are assessed. When a binary representation is applied to DMR data with and without covariates, there is an assumption that all DMRs that change will also have the same effect size changes and would be equally important for motif discovery. In contrast, accounting for the continuous range of differential methylation and effect size changes with covariates can provide a more accurate representation of the potential transcription factors involved with differential methylation at specific loci. With the SArKS method, we found numerous motifs and transcription factor binding site predictions that change when including postnatal maternal licking/grooming as a covariate, as we would have expected based on the DMR analysis. This was also the case with the MEME Suite method, and the changes were sometimes even more pronounced, possibly due to the binary representation of the input sequences as described above.

A consistent theme throughout our analyses was that the genes and transcription factor binding site predictions associated with prenatal bisphenol exposure and postnatal maternal licking/grooming as well as their interactions were either related to estrogen receptor signaling or were estrogen-responsive genes. In prior transcriptome analyses in the PND10 prefrontal cortex and amygdala, we found consistently strong and significant enrichment of ESR1 in the Transcription Factor Protein-Protein Interactions module in Enrichr [26]. This was also the case in the PND10 MPOA, but we also found significant (though weaker) enrichment of ESR1 in the ENCODE and ChEA Consensus TFs from ChIP-X module. These findings are consistent with our hypothesis that the interactions between prenatal bisphenol exposure and postnatal maternal licking/grooming converge on estrogen receptor signaling, with some brain region-specific differences, and that we are primarily finding secondary effects of estrogen receptor signaling changes by PND10.

Finally, we examined the mitigating effect of licking-like tactile stimulation on prenatal bisphenol-exposed pups by directly manipulating the levels of tactile stimulation received between siblings within a litter. This within-litter study design allowed us to control for other possible within-litter effects and partially control for pup genotype. We focused on female pups and the 150 μg/kg bisphenol dose based on the prior tag-seq data. While we found widespread reductions in DMRs throughout the genome between the prenatal bisphenol-exposed pups and Corn Oil controls in the stimulated siblings, the effect sizes for individual DMRs were generally small since we only observed a few interactive effects in our DMR analyses. However, the DMRs that did show interactions between prenatal treatment and postnatal tactile stimulation condition were within estrogen-responsive genes. We also found mitigating effects of supplemental tactile stimulation on ESRRG binding to ESRRG-responsive genes—prenatal bisphenol-exposed pups had reduced ESRRG binding which was normalized in the stimulated siblings. The reduction in ESRRG binding might be associated with hypermethylation of the gene in most cases. Like the DMR analysis, the effect sizes were relatively small, and we found only one statistically significant interaction between 150 μg/kg Mixed BP exposure and postnatal tactile stimulation condition with Foxp1. Nevertheless, the effects of supplemental tactile stimulation on ESRRG binding were consistent across all the ESRRG-responsive genes we tested, even if the differences were not always statistically significant. Interestingly, the same patterns of binding enrichment occurred regardless of the DMR results, which reflects some discrepancies between ESRRG binding and DNA methylation changes. This does not appear to be strongly related to the distance between the DMR and transcription factor binding site. Like the gene expression data, ESRRG binding is a transient event while DNA methylation is relatively stable, and there is likely a delay between ESRRG binding and DNA methylation changes. The discrepancy also illustrates that the links between transcription factor binding and DNA methylation changes are not always straightforward at each gene loci. This may also explain why we did not find significant enrichment of EREs or ESRRG binding motifs in the differentially methylated transcription factor binding site analyses. Another important discrepancy is that we found reductions in ESRRG binding in nonstimulated pups with prenatal bisphenol exposure while we also found increases in *Esrrg* expression in prenatal-bisphenol pups with lower levels of postnatal licking/grooming received. While it has been shown that acute BPA exposure increases ESRRG activity, there may be changes in ESRRG activity over longer exposure periods that reflect an adaption to the exposure. One prior study in rare minnows has found opposing ESRRG activity changes with short-term and long-term BPA exposure [60]. Prior work has also found discrepancies between expression of steroid hormone receptors and the binding of those steroid hormone receptors to gene loci [37], so gene expression data may not be predictive of DNA binding activity. Indeed, we found in our study that *Esrrg* expression had both positive and negative connectivity to the co-expressed gene modules constructed in our WGCNA analysis, demonstrating that *Esrrg* expression is also negatively correlated with the expression of many other genes. With this information in mind, it is still possible that ESR1 binding activity may account for some of the mitigating effects of postnatal licking/grooming or supplemental tactile stimulation observed in this study. We did not find differences in ESRRG binding to the ERE in the Septin9 gene, even though enrichment was overall higher than the other ESRRG-responsive genes. While it is currently unknown whether Septin9 is an estrogen-responsive gene, which is a limitation in this study, we did identify a few known ESR1-responsive genes in our DMR analyses (including Ampd3) that could be studied more deeply in future work.

One other notable limitation of this study was while we found consistent interactions between postnatal licking/grooming or supplemental tactile stimulation and the 150 μg/kg bisphenol dose, this dosage is not considered environmentally relevant and is generally higher than human exposure levels. However, bisphenol exposure follows a non-monotonic dose response curve [61,62], meaning that very high levels and very low levels will exert biological effects. As an example, in our study, the 50 μg/kg Mixed BP and BPA-only dose did not facilitate any increases of *Esrrg* expression in female pups **(Supplementary Figure S1A**); however, *Esrrg* expression may change with doses lower than 50 μg/kg, which would be environmentally relevant. Consistent with this logic, our lab’s previous work on prenatal BPA exposure found that the 2 μg/kg and 20 μg/kg doses alter *Esrrg* expression in the prefrontal cortex and hypothalamus [19]. While the dosages used in the current study were informative for discovering potential biological mechanisms, additional studies using lower doses are needed to ensure the translational potential of our current findings. Finally, we do not have any data on how maternal licking/grooming might mitigate the impacts of prenatal bisphenol exposure on MPOA-dependent behaviors later in life, such as social behavior and maternal care provisioning to the next generation. However, given that there is already evidence that both prenatal bisphenol exposure and postnatal maternal care can impact these later-life phenotypes [63–69], it is likely that interactions between these two early-life environmental exposures exist.

## 4. Conclusions

Overall, we found that postnatal maternal licking/grooming can mitigate the impacts of prenatal bisphenol exposure on *Esrrg* expression and co-expressed gene modules in the developing MPOA in a dose-dependent and sex-specific manner. This appears to correspond to changes in DNA methylation, which can persist beyond both transient environmental exposures and impact neurodevelopmental trajectories and behavioral phenotype. We also found mitigating effects of licking-like tactile stimulation on DNA methylation and ESRRG binding activity in the developing MPOA of female offspring, which indicates that supplemental tactile stimulation could be implemented as an effective intervention against the adverse neurodevelopmental impacts of prenatal bisphenol exposure. Overall, these results have identified one potential avenue to mitigate the effects of prenatal bisphenol exposure and improve health and well-being in human populations, which warrants further exploration in future studies.

## 5. Materials and Methods

**Supplementary Table S17.** displays all the code files and data files used for the bioinformatic analyses, as well as the information from all samples used for analyses.

### 5.1 Husbandry and breeding

This study examined the effects of naturally occurring variations in licking/grooming (Maternal LG; Figure 1A) and experimentally induced increases in licking-like tactile stimulation (Tactile LG; Figure 1B). All animal procedures were approved by the Institutional Animal Care and Use Committee at the University of Texas at Austin and conformed to the guidelines of the American Association for Laboratory Animal Science. The rat pups used for the Maternal LG analyses were part of a larger cohort that has been described in previous publications [26,59]. The rat pups used for the Tactile LG analyses have not been previously published. Briefly, breeder female and male Long-Evans rats (Charles River) were housed in same-sex pairs on a 12:12 hour inverse light-dark cycle with ad libitum access to standard chow diet (#5LL2, Lab Diet) and water. To limit external exposure to bisphenols and other xenoestrogens, all rats were provided with glass water bottles, polysulfone cages, and aspen wood shavings for bedding material.

### 5.2 Gestational bisphenol administration

Adult male and female rats were screened for breeding receptivity by pairing one male with one or two females and observing mounting behavior for the male and lordosis for the female(s). From gestational day 8 until parturition, cage-pairs of dams were randomly assigned to receive control Corn Oil or one of the bisphenol (BP) exposure treatments. The Maternal LG analyses included 50 μg/kg BPA, 50 μg/kg Mixed BP, or 150 μg/kg Mixed BP. Based on the observed effects on transcription of the Maternal LG doses (**Figures 2 & 3**), the Tactile LG analyses included 150 μg/kg BPA or 150 μg/kg Mixed BP. The timing of BP dosing was selected based on prior work demonstrating that gestation day 8 to parturition encompasses the start of brain sexual differentiation and when the expression of estrogen receptors is apparent in the fetal brain [70]. Stock solutions of bisphenol A (BPA; #B0494, TCI, ≥ 99 %), bisphenol S (BPS; #A17342, Alpha Aesar, ≥ 99 %), or bisphenol F (BPF; #A11471, Alfa Aesar, ≥ 98 %) were dissolved in corn oil (#405435000, Acros Organics). Equal parts BPA, BPS, and BPF were used for both Mixed BP treatment groups. Experimenters involved with the administration of treatments were blinded to the treatment groups. On the day of parturition (postnatal day 0; PND0), offspring sex was determined using relative anogenital distance and litters culled to 5-6 male and 5-6 female pups per litter.

### 5.3 Postnatal maternal care observations and quantification

For the Maternal LG analyses, home cage maternal behavior was recorded for one hour per day starting one hour after lights-off from PND1-10 using Raspberry Pi 3B+ minicomputers. Maternal behavior was scored by either manual coding or through the AMBER pipeline [71]. All postnatal maternal care measures were normalized to seconds of observed behavior per day and then scaled and centered where the average is 0 and standard deviation is 1 for all downstream bioinformatic analyses. Based on prior analyses [26,59] we used licking/grooming and nest attendance from PND1-5 as independent variables or covariates for the current study. Maternal behavior measures used in the current study did not vary between prenatal bisphenol exposure groups [26]. Following the maternal care video recordings at PND10, the brains from one male and one female pup were collected and flash-frozen in hexanes on dry ice.

### 5.4 Postnatal supplemental licking-like tactile stimulation administration

The supplemental tactile stimulation procedure used in the Tactile LG analyses was derived from a previous study [37] with some modifications. Each day from PND2-10, litters were separated from the dam for approximately 25 minutes per day and transported to a separate room starting one hour after lights-off. Whole litters were transported into small cages lined with aspen wood shavings that were warmed to thermoneutral conditions (33-35°C) using a heating pad (#RT-0520, Kent Scientific). During the daily 25-minute separation, 2-3 male pups and 2-3 female rat pups within a litter received licking-like tactile stimulation with a camel hair paintbrush (Craftsmart) for 15 minutes (‘stimulated’ condition) while the remaining pups were left undisturbed (‘nonstimulated’ condition). Pups received the supplemental tactile stimulation on the dorsal region of their body at a rate of approximately two strokes per second. The same pups received supplemental tactile stimulation each day. Pups of the same sex were weighed together daily and the interscapular temperature was periodically monitored with an infrared thermometer (#33501-416, VWR) throughout the separation to ensure the pups did not experience hypothermia. From PND2-9, rat pups that were assigned to the supplemental tactile stimulation condition were marked using odourless and tasteless food colouring (McCormick), as described in previous work [37,72]. Immediately after the final tactile stimulation administration, the brains from all pups were collected and flash-frozen in hexanes on dry ice. One stimulated and one nonstimulated same-sex sibling were simultaneously euthanized at a time. Only female pup brains were used for downstream analyses in cohort 2.

### 5.5 RNA extraction and 3’ tag sequencing (Tag-seq)

Power analyses were done using the R package PROPER for RNA-seq data with the Gilad dataset (medium biological variation) and 25 data simulations to calculate effect size.Analyzing 6 biological samples per group allowed 72% power to correctly identify differentially expressed genes with an FDR value < 0.05 and log fold change of at least 0.5.

All brains were stored at −80°C until cryosectioning. PND10 brains (Maternal LG analyses; n = 6 per sex per prenatal treatment group) were cryosectioned in 50 μm slices using a ThermoFisher Scientific CryoStar NX50 cryostat or a Leica CM3050S cryostat. The medial preoptic area (MPOA; 0.20 to −0.60 mm Bregma) was microdissected using an atlas for the developing rat brain [73] and a supplementary atlas for the PND10 rat brain [74]. RNA was extracted using the MagMAX FFPE DNA/RNA Ultra Kit (#A31881, ThermoFisher Scientific). For unfixed frozen tissue, 210 μl of protease solution was pipetted onto each MPOA sample and allowed to incubate at 55°C at 900 rpm for at least one hour. The homogenate was immediately processed for RNA and DNA extraction using the Kingfisher Flex System following the manufacturer’s instructions. RNA quantity was assessed with the Quant-iT RNA Assay kit (#Q33140, ThermoFisher Scientific) and RNA quality was assessed using the RNA 6000 Pico Assay kit (#5067-1513, Agilent Technologies). All RNA samples were diluted to 20 ng/μl before submission for library preparation and tag-seq [75,76] at the Genome Sequence and Analysis Facility at the University of Texas at Austin. Reads were sequenced on the NovaSeq S1 (100 bp single-end reads).

### 5.6 DNA extraction and oxidative reduced representation bisulfite sequencing (oxRRBS)

For the Maternal LG analyses, DNA was extracted using the MagMAX FFPE DNA/RNA Ultra Kit (#A31881, ThermoFisher Scientific). A subset of samples (two 150 μg/kg Mixed BP females, one 150 μg/kg Mixed BP male) did not contain sufficient DNA for library preparation for oxRRBS. Another subset of samples (one 50 μg/kg BPA male, one 150 μg/kg Mixed BP female, one Corn Oil female) likely had substantial anterior pituitary gland contamination based on the tag-seq results (described in the tag-seq preprocessing section) and were not processed for oxRRBS. To maintain a sample size of 6 per sex per prenatal treatment group, DNA was extracted from additional MPOA samples using the MagMAX DNA Multi-Sample Ultra 2.0 Kit following the manufacturer’s instructions for tissue samples (#A45721, ThermoFisher Scientific). For the Tactile LG analyses, DNA was extracted using the MagMAX DNA Multi-Sample Ultra 2.0 Kit following the manufacturer’s instructions for tissue samples. DNA quantity was assessed with the Quantifluor dsDNA system (#E2670, Promega).

Library preparation for oxRRBS was done using the Ovation RRBS Methyl-Seq System with the TrueMethyl® oxBS module (Tecan) using 55 ng of input DNA. For the Maternal LG analyses, between 12-20 samples were processed per batch of library preparation with a total of 5 batches, balanced by prenatal treatment group and sex. For the Tactile LG analyses, 12 samples were processed per batch of library preparation with a total of 3 batches, balanced by prenatal treatment group and postnatal tactile stimulation condition. The oxidation of DNA prior to bisulfite conversion converts 5hmC to 5fC and allows for discrimination between 5mC and 5hmC at single base-pair resolution. For this study, oxidative bisulfite conversion was used to remove 5hmC and detect only 5mC in the samples. Final libraries were quality-checked using the Agilent Technologies High Sensitivity DNA Kit. Five libraries from the Maternal LG analyses (one 50 μg/kg BPA female, one 50 μg/kg Mixed BP male, one 50 μg/kg Mixed BP female, and two 150 μg/kg Mixed BP females) and one library from the Tactile LG analyses (one 150 μg/kg Mixed BP female with tactile stimulation) were removed due to insufficient final yield. The remaining libraries were pooled with 8-9 libraries per pool for the Maternal LG analyses (balanced by prenatal treatment group and sex) and 5-6 libraries per pool for Tthe actile LG analyses (balanced by prenatal treatment group and postnatal tactile stimulation condition). Library pools underwent an additional Ampure XP bead clean-up and were submitted for sequencing to the Genome Sequence and Analysis Facility at the University of Texas at Austin. Reads were sequenced on the NovaSeq SP (150 bp paired-end reads).

### 5.7 Chromatin Immunoprecipitation (ChIP) for ESRRG

For the Tactile LG analyses, five MPOA samples per prenatal treatment group and postnatal tactile stimulation condition were processed for ChIP based on previously used protocols [37,77]. Five batches were conducted with one MPOA sample per prenatal treatment group and postnatal tactile stimulation condition per batch. MPOA samples were cross-linked with 1% formaldehyde (Sigma-Aldrich) for 10 minutes at 24°C. The samples were quenched with 1.25 M glycine and left on a rotating platform (50 rpm) at room temperature for 5 minutes, centrifuged at 21,000g for 30 seconds and washed five times on ice with ice-cold 1x PBS and a protease inhibitor cocktail (Thermofisher) dissolved in 1x PBS. MPOA tissue was then homogenized with a SDS lysis buffer (0.25 M sucrose, 60 mM KCl, 15 mM NaCl, 10 mM MES (pH 6.5), 5 mM MgCl_2_, 0.5% Triton X-100) and centrifuged at 4,500g at 4°C to pellet the cell nuclei. The SDS lysis buffer was removed and a salt buffer (50 mM NaCl, 10 mM PIPES (pH 6.8), 5 mM MgCl_2_, 1 mM CaCl_2_) was added prior to sonication (low setting; three times for 10 seconds on, 30 seconds off; Diagenode Bioruptor® Plus sonication device). The samples were incubated with 150 units of micrococcal nuclease (#10011S, Cell Signaling) at 37°C for 35 minutes before quenching with 5 µl 0.5 M EDTA and placed on ice. SDS (10%) was added to each sample before being centrifuged at 21,000g for 5 minutes, aliquoted and diluted 4x with a ChIP dilution buffer. Each ChIP aliquot contained 20 µl of Protein G magnetic beads (#16-662, Millipore) and 10 µg ESRRG antibody (#PPH681200, R&D Systems; [78]) and was incubated on a rotating platform (50 rpm) at 4°C overnight. The beads were then pelleted using a magnetic separator and washed with ice-cold low-salt, high-salt, LiCl (#20-156, Sigma) and Tris-EDTA buffers. Cross-links were reversed for ChIP aliquots and input chromatin samples using 10 µg of proteinase K in Tris-EDTA buffer (with 1% SDS) at 65°C for at least 2 hours before purification using the Qiaquick PCR purification kit (Qiagen).

The ESRRG DNA binding motifs for most candidate genes tested (Eno4, Foxp1, Ncor2, Ppp1r3b) were identified by the rat homologous sequences from previous literature using mouse neuron ChIP-seq data [55]. The ERE from Septin9 was identified by aligning the consensus palindromic motif (5′-GGTCAnnnTGACC-3′) to the rat Septin9 genetic sequence. Primers for all candidate genes were created with Primer-BLAST software (**Supplementary Table S18**). Enrichment was measured for each gene locus using quantitative PCR (Fast SYBR Green technology) for the ChIP DNA and input chromatin samples with technical triplicates. The enrichment for ESRRG was normalized and calculated as relative percentages of input chromatin. Due to the relatively low enrichment of ESRRG overall, technical replicates were removed from enrichment calculations if there was no amplification or notable primer dimer found in the melt curve analysis. If there was no amplification or notable primer dimer was found in the melt curve analysis in all technical replicates, the sample was considered to have 0% enrichment. Statistical analyses were performed using R version 4.1.3 (R Core Team, 2023). Linear mixed models were performed using the lme4 R package using batch and antibody tube used as random factors for all analyses. To examine main effects of prenatal bisphenol treatment and interactions with postnatal tactile stimulation condition, 2×2 linear mixed models were conducted. Comparisons between stimulated and nonstimulated groups within each prenatal treatment group were also done to examine the effects of supplemental tactile stimulation on ESRRG enrichment. All effects were reported as statistically significant if p ≤ 0.05 and marginally significant if p ≤ 0.10.

### 5.8 Pipeline and Analyses: Tag-seq

#### 5.8.1 Preprocessing

The tag-seq pipeline overview can be found in **Figure 1C** and all data and code for the differential gene expression and weighted gene co-expression network analysis can be found at https://github.com/SLauby/bisphenols_maternalcare_epigenome. The analysis pipeline followed previously published tag-seq analyses [26]. Adaptor sequences, poly-A tails, and low-quality reads were trimmed (https://github.com/z0on/tag-based_RNAseq) followed by cutadapt (version 1.18). Trimmed reads were mapped to the rat genome (mRatBN7.2; RefSeq) using STAR (version 2.7.3a) with the default settings and sorted using samtools (version 1.10). About 90% of reads were uniquely mapped to the rat genome for all samples. Counts for each gene were calculated for each sample using the bedtools (version 2.27.1) multicov function. Three samples (one 50 μg/kg BPA male, one 150 μg/kg Mixed BP female, one Corn Oil female) had unusually high counts (over 100x of the other samples) for anterior pituitary gland-specific genes, including *Pomc, Gnrhr,* and *Prl*. These samples were removed from the dataset prior to all downstream analyses.

#### 5.8.2 Differential gene expression analysis

For all downstream tag-seq analyses, comparisons between prenatal treatment groups and interactions with prenatal treatment group and postnatal maternal care were always done relative to the Corn Oil group. To examine differential expression of individual genes, including all estrogen and estrogen-related receptors, gene count data was normalized and analyzed for differential expression using the DESeq2 R package (version 1.34.0), separated by sex. Genes with more than 5 counts for more than 6 samples were analyzed. Prenatal treatment and postnatal maternal care measures (licking/grooming, nest attendance) from PND1-5 were used as independent variables. The estrogen and estrogen-related receptors (*Esr1, Esr2, Esrra, Esrrb, Esrrg*) were specifically examined for significant interactions between each prenatal treatment group and licking/grooming. Other potential differentially expressed genes (DEGs) were identified with a false discovery rate (FDR) alpha ≤ 0.10 and ≤ 0.05 with Benjamani-Hochburg correction. All effects were reported as statistically significant if p ≤ 0.05 and marginally significant if p ≤ 0.10.

#### 5.8.3 Weighted gene co-expression network analysis (WGCNA)

To examine differential expression of co-expressing gene profiles, gene networks were created with the WGCNA R package (version 1.72-1) using normalized and log-transformed gene count data, split by sex. Genes with very low counts and variation were filtered out using the goodSamplesGenes function in R (verbosity = 3). WGCNA calculates Pearson correlations of count data between every gene for network construction and clusters highly correlated genes into eigengene modules (at least 30 genes per module). A soft power threshold of 8 was used for network construction. We excluded modules with eigengene values that were characterized by one extreme outlier (> 0.90) and low variation following outlier removal (standard deviation < 0.10) from further analyses. The eigengene values from the remaining modules were analyzed using the lm function in R. Prenatal treatment and the postnatal maternal care measures (licking/grooming, nest attendance) from PND1-5 were used as independent variables. For any eigengene module that showed statistically significant interactions, the gene sets were extracted and analyzed using Enrichr [79,80] using the full gene list used for network construction as the background (https://maayanlab.cloud/Enrichr/). To examine if the gene sets were enriched in estrogen-responsive genes, ESR1 was examined in the ENCODE and ChEA Consensus TFs from ChIP-X module. To examine if the gene sets were enriched in genes that are known to interact with estrogen receptors, ESR1 was examined in the Transcription Factor Protein-Protein Interaction module. To examine the degree of connectivity with the co-expressed gene modules with specific estrogen receptors, the kME was calculated for all estrogen and estrogen-related receptors in the WGCNA package. The kME determines how closely a gene of interest is related to a particular module and is represented as a positive or negative percentage of all genes that have highly correlated expression to the gene of interest. Negative values indicate inverse relationships. All effects were reported as statistically significant if p ≤ 0.05 and marginally significant if p ≤ 0.10.

### 5.9 Pipeline and Analyses: oxRRBS

#### 5.9.1 Preprocessing

The oxRRBS pipeline overview can be found in **Figure 1C** and all code for the differentially methylated region analyses can be found at https://github.com/SLauby/bisphenols_maternalcare_epigenome. The bismark-processed DNA methylation datasets can be accessed in Zenodo (https://doi.org/10.5281/zenodo.17273659). Adaptor sequences and low-quality reads (q < 30) were trimmed using trim galore (version 0.6.10) and cutadapt (version 4.6). Diversity sequences and reads without the MspI site (CGG) were removed using a custom python script provided by the manufacturer of the oxRRBS kit (https://github.com/nugentechnologies/NuMetRRBS). Paired-end reads were mapped to the rat genome (mRatBN7.2; RefSeq) using bismark (version 0.22.3) and bowtie2 (version 2.5.2) with the default settings. About 75% of reads were uniquely mapped to the rat genome for all samples. PCR duplicates were removed based on the genomic coordinates of the reads and a 6-bp unique molecular identifier found in the read index using a custom python script provided by the manufacturer of the oxRRBS kit (https://github.com/tecangenomics/nudup). For the Maternal LG analyses, there was an average of about 8.5 million uniquely aligned, de-duplicated reads per sample. For Tthe actile LG analyses, there was an average of about 6.5 million uniquely aligned, de-duplicated reads per sample. All samples were analyzed for DNA methylation levels using the bismark methylation extractor. Bisulfite conversion rate was > 99% for all samples.

#### 5.9.2 Differentially methylated region (DMR) analysis

For all downstream oxRRBS analyses, pairwise comparisons between prenatal treatment groups were always done relative to the Corn Oil group. For the Tactile LG analyses, pairwise comparisons between prenatal treatment groups were done for pups within the same postnatal tactile stimulation condition and pairwise comparisons between the postnatal tactile stimulation conditions were done for pups within the same prenatal treatment group. Differentially methylated regions (500 bp) between prenatal treatment groups were examined using Methylkit (version 1.26.0), separated by sex for the Maternal LG analyses. Regions were included in the analysis if there were over 10x reads for at least 4 samples per prenatal treatment group (and postnatal tactile stimulation condition for the Tactile LG analyses). Batch of library preparation was used as a covariate in all analyses. DMRs were identified with an FDR q-value ≤ 0.05 using the SLIM method and had > 5% overall methylation difference between groups. DMR analyses are currently limited to pairwise comparisons, so an alternative approach was undertaken to examine potential interactions between prenatal bisphenol exposure and postnatal maternal licking/grooming or tactile stimulation. For the Maternal LG analyses, each comparison between the prenatal treatments groups and Corn Oil group was done including and excluding postnatal maternal licking/grooming as a covariate and the degree of overlap in DMRs was identified. DMRs that were lost or added when including postnatal maternal licking/grooming as a covariate in the analysis were considered to be influenced by postnatal maternal licking/grooming. More specifically, DMRs that were added were assumed to be due to interactions between prenatal bisphenol exposure and postnatal maternal licking/grooming. For the Tactile LG analyses, each comparison between the prenatal treatments groups and Corn Oil group was done including and excluding postnatal tactile stimulation condition as a covariate.

#### 5.9.3 Differentially methylated motif and transcription factor binding site analysis: Suffix Array Kernel Smoothing (SArKS)

To adapt the SArKS method to RRBS data, genomic regions with methylation data were created as input sequences for the SArKS algorithm to identify overrepresented sequence motifs. The code for creating those input sequences can be found at https://github.com/denniscwylie/prenatal_bp_x_lg_epigenome_analysis. For each individual sample, observed methylation sites were partitioned into disjointed genomic ranges separated by at least 51 base pairs. The bedtools merge command was then applied (using the bedtoolsr::bt.merge function in a custom R script) to combine the resulting partitioned range sets from each individual sample into a single set of partitioned genomic ranges in which every individual sample range is contained within one of the merged range intervals. The merged set of genomic ranges were then trimmed using the bedtools coverage command together with a custom R script, only retaining positions which were covered by one of the individual sample-partitioned ranges for at least 25 (out of 43 total) of the individual samples analyzed. For each sample, a local methylation rate was assigned to each genomic position within the merged set of (trimmed) genomic ranges defined above. Each individual base position within each of these partitioned regions is assigned a local methylation rate equal to the estimated methylation percentage for the nearest site observed methylation site (within 50 bp). If the nearest site observed methylation site is > 50 bp, the local methylation rate is considered undefined for the sample. Finally, a single average methylation level for each sample was then computed for each of the merged (and trimmed) genomic ranges as the mean of the assigned individual base methylation rates over all positions within the range (omitting any positions with undefined local methylation rates).

For each genomic range R in the merged (and trimmed) set, a linear model was incorporated to model the arcsin-transformed range-averaged methylation rates as the dependent variable and the library preparation batch covariate and prenatal treatment group as independent predictor variables. Each comparison was done including and excluding postnatal maternal licking/grooming as a covariate. In this way, an estimated model coefficient was obtained for each combination of genomic range R and prenatal bisphenol treatment group to the Corn Oil group with and without postnatal maternal licking/grooming as a covariate. For each prenatal bisphenol treatment group comparison, SArKS (https://github.com/denniscwylie/sarks; [81]) was used to identify overrepresented sequence motifs found within the various merged genomic ranges whose presence within genomic range R correlates with more extreme (either higher or lower, depending on which direction the analysis was run) estimated model coefficients for each prenatal treatment group comparison associated with the region R. Step-by-step instructions and pseudocode for SArKS can be found at https://bioconductor.posit.co/packages/3.23/bioc/vignettes/sarks/inst/doc/sarks-vignette.pdf.

#### 5.9.4 Differentially methylated motif and transcription factor binding site analysis: MEME Suite

For the Maternal LG analyses, DMRs that were found using Methylkit were used to identify overrepresented sequence motifs with MEME Suite (Version 5.5.7; [82]) using previously published parameters [83] with some modifications. To make the analysis more comparable to SArKS, we included regions that had differential methylation with a nominal p-value ≤ 0.05 and had > 0% overall methylation difference between groups. In addition, MEME was used in discriminative mode using a set of sequences that were not differentially methylated between the prenatal treatment groups and Corn Oil group.

For both SArKS and MEME Suite methods, overrepresented sequence motifs were examined for potential transcription factor binding sites using the JASPAR 2024 CORE (non-redundant) database for vertebrates [84]. E-values ≤ 0.05 were considered statistically significant.

### 5.10 Multi-omic factor analysis (MOFA) of tag-seq and oxRRBS datasets

Directly comparing gene expression differences and DNA methylation modifications can be challenging when the analysis pipeline for each measure is not integrated. To examine the interactions between prenatal bisphenol exposure and maternal care on the epigenome in a more holistic manner, a MOFA was performed using the MOFA2 package in R (version 1.10.0), separated by sex. Multi-omic analyses can only be performed if the tag-seq and oxRRBS datasets were derived from the same MPOA samples; therefore, the sample sizes were smaller than either the tag-seq or oxRRBS analyses (n = 3-6 per sex per prenatal treatment group). Gene count data were normalized with DESeq2. DNA methylome data were restricted to one 500 bp region per 2000 bp window to reduce the influence of the high covariance structure of DNA methylation in proximal regions. Proportion of DNA methylation (p) was then log-transformed, similar to M-values in microarray datasets:

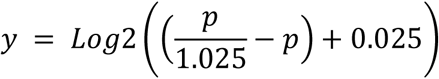

The gene count dataset was filtered to only include the top 5,000 most variable genes and the DNA methylome dataset was filtered to only include the top 10,000 most variable regions. MOFA2 was performed using the default options, except that the scale views option was set to TRUE, convergence mode was set to slow, and the number of factors was reduced to 5. The weights from each sample and factor were extracted and analyzed using the lm function in R. Prenatal treatment and the postnatal maternal care measures (licking/grooming, nest attendance) from PND1-5 were used as independent variables. For any factor that showed statistically significant main effects or interactions, the top genes and regions by weight (250 positive weights and 250 negative weights) were extracted. The most proximal gene was determined from each region (if within the promoter or gene body) from the oxRRBS dataset and examined for any overlap with the top genes from the tag-seq dataset. The top genes (removing the uncharacterized genes) from the tag-seq and oxRRBS datasets were separately analyzed with Enrichr. Enrichment for ESR1 was examined in the list of ENCODE and ChEA Consensus TFs from ChIP-X and Transcription Factor Protein-Protein Interaction modules. All effects were reported as statistically significant if p ≤ 0.05 and marginally significant if p ≤ 0.10.

## Supporting information

Supplementary Results

## 6. Acknowledgements

We thank Isha Agarwal, Taylor Hite, and Madeline Severson for assisting the prenatal bisphenol administration for the Maternal LG analyses, and Jean Botani for assisting the prenatal bisphenol administration and postnatal supplemental tactile stimulation manipulations for the Tactile LG analyses.

