## Supplementary Figures for "Postnatal maternal care normalizes the hypothalamic DNA methylome following prenatal bisphenol exposure"

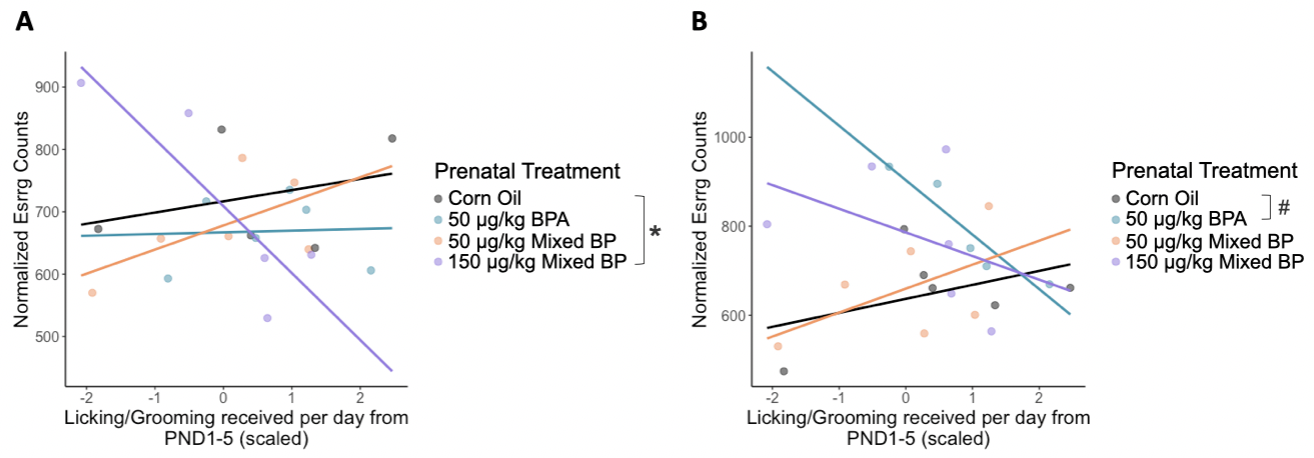

**Supplementary Fig S1.** Sex-specific interaction between prenatal bisphenol exposure and postnatal maternal licking/grooming on *Esrrg* expression in the medial preoptic area. A significant interaction between the 150 μg/kg Mixed BP exposure group and maternal licking/grooming was found in female pups (**A**) but not male pups (**B**). In females from the 150 μg/kg Mixed BP group, maternal licking/grooming was negatively correlated with Esrrg expression. Scatterplots are displayed with linear regression lines for each prenatal treatment group. * p < 0.05 interaction between prenatal treatment and postnatal maternal care; # p < 0.10 interaction between prenatal treatment and postnatal maternal care

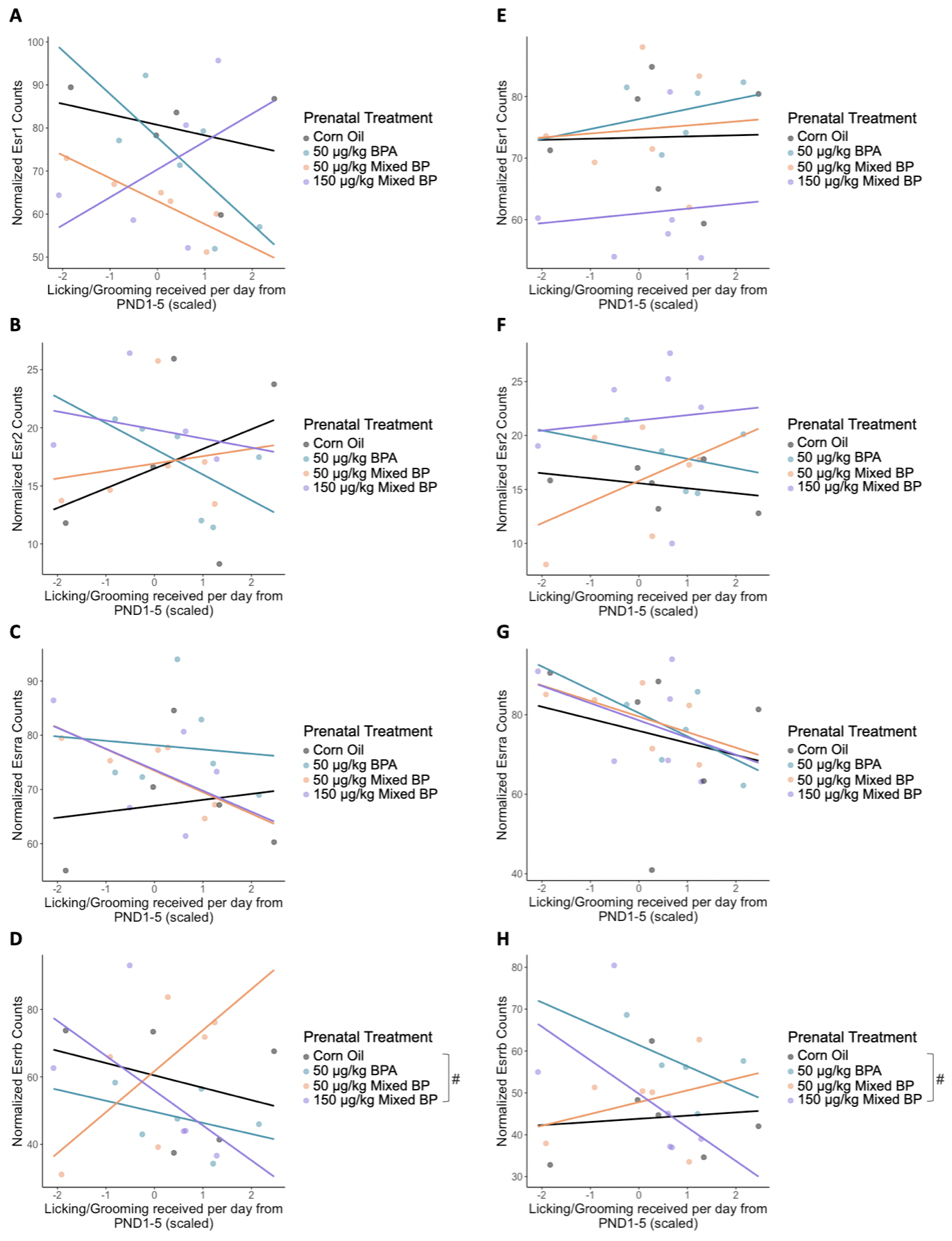

**Supplementary Fig S2.** No significant interactions between prenatal bisphenol exposure and postnatal maternal licking/grooming on expression levels of other estrogen and estrogen-related receptors in the medial preoptic area. This included *Esr1, Esr2, Esrra*, and *Esrrb* in female (**A-D**) and male (**E-H**) pups. Scatterplots are displayed with linear regression lines for each prenatal treatment group. # p < 0.10 interaction between prenatal treatment and postnatal maternal care

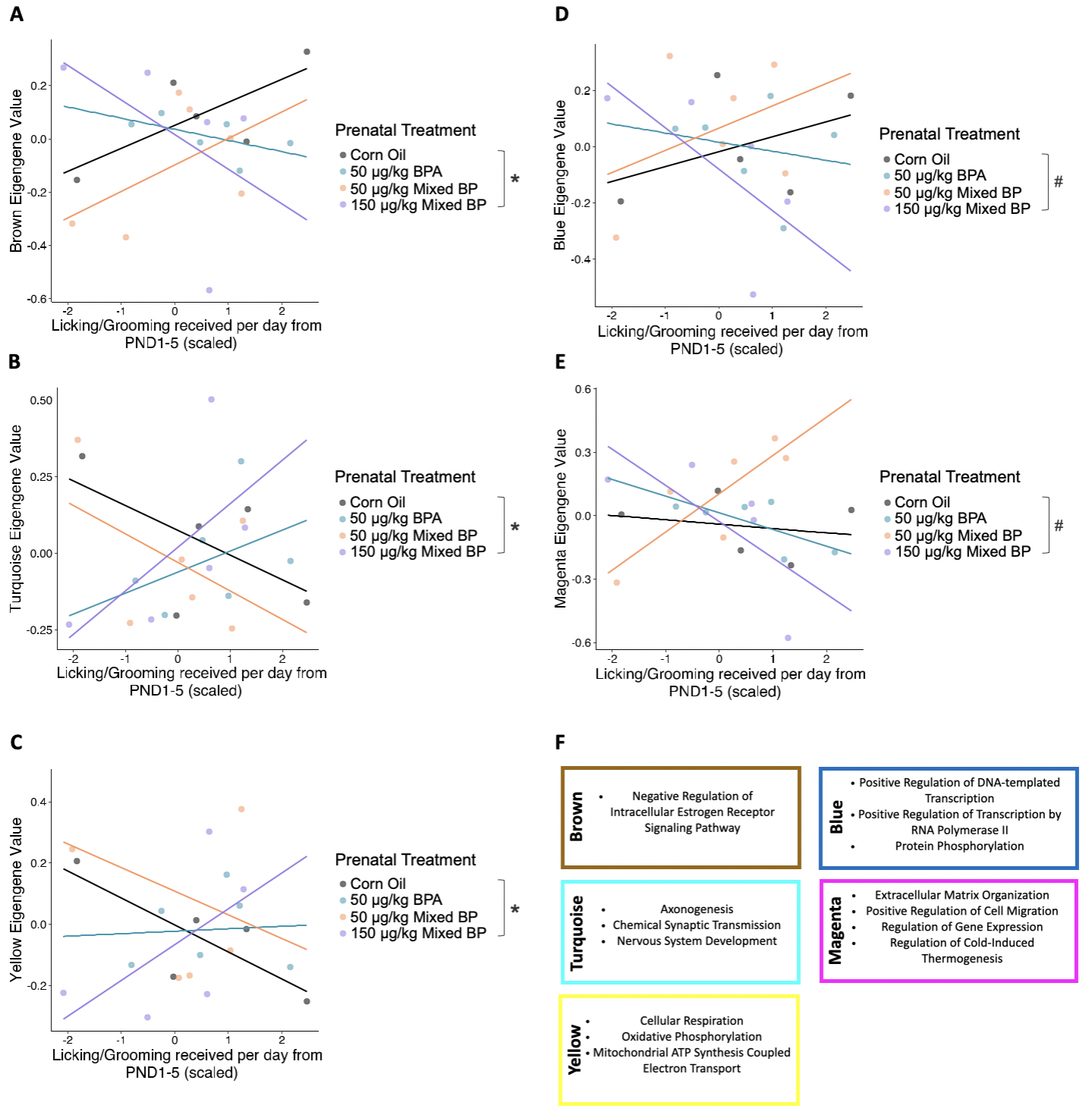

**Supplementary Fig S3.** Interactions between prenatal bisphenol exposure and postnatal maternal licking/grooming on co-expressed gene modules in the medial preoptic area of female pups. Significant interactions between the 150 μg/kg Mixed BP exposure group and maternal licking/grooming were found in the Brown (**A**), Turquoise (**B**), and Yellow (**C**) eigengene modules. Marginal interactions were found in the Blue (**D**) and Magenta (**E**) eigengene values. In females from the 150 μg/kg Mixed BP group, maternal licking/grooming was negatively correlated with the Brown, Blue, and Magenta eigengene values and positively correlated with the Turquoise and Yellow eigengene values. Top GO terms (**F**) from these modules were related to estrogen receptor signaling, regulation of gene expression, neurodevelopment, neurotransmission, and cellular metabolism. Scatterplots are displayed with linear regression lines for each prenatal treatment group. * p < 0.05 interaction between prenatal treatment and postnatal maternal care; # p < 0.10 interaction between prenatal treatment and postnatal maternal care

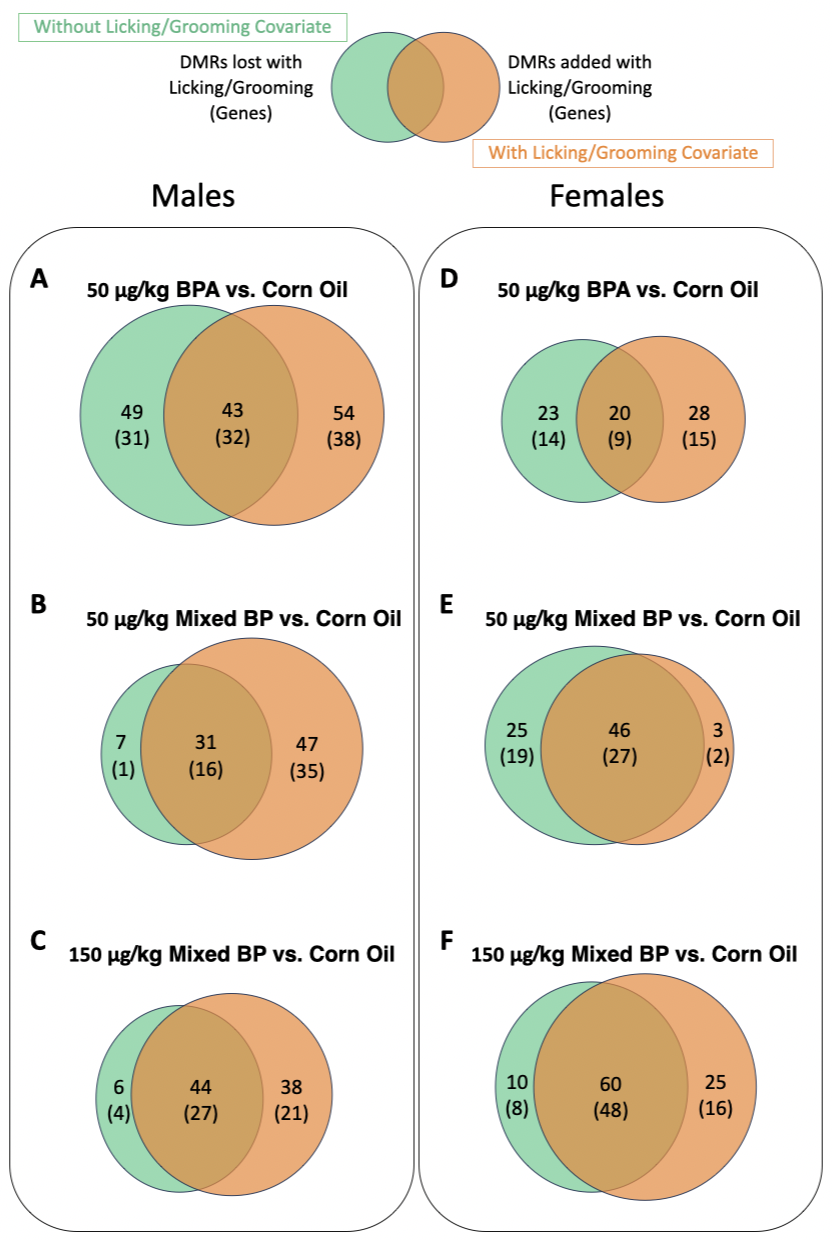

**Supplementary Fig S4.** Influence of postnatal maternal licking/grooming on differentially methylated regions (DMRs) of prenatal bisphenol-exposed male and female pups. Including maternal licking/grooming as a covariate altered DMRs in all prenatal treatment groups in both male (**A-C**) and female (**D-F**) pups. For all comparisons except for the females in the 50 μg/kg Mixed BP group, more DMRs were added than lost when adding licking/grooming as a covariate. Venn diagrams are displayed with the number of DMRs and annotated genes when excluding (green) and including (orange) maternal licking/grooming as a covariate. DMRs that do not overlap are considered to be influenced by postnatal maternal licking/grooming.

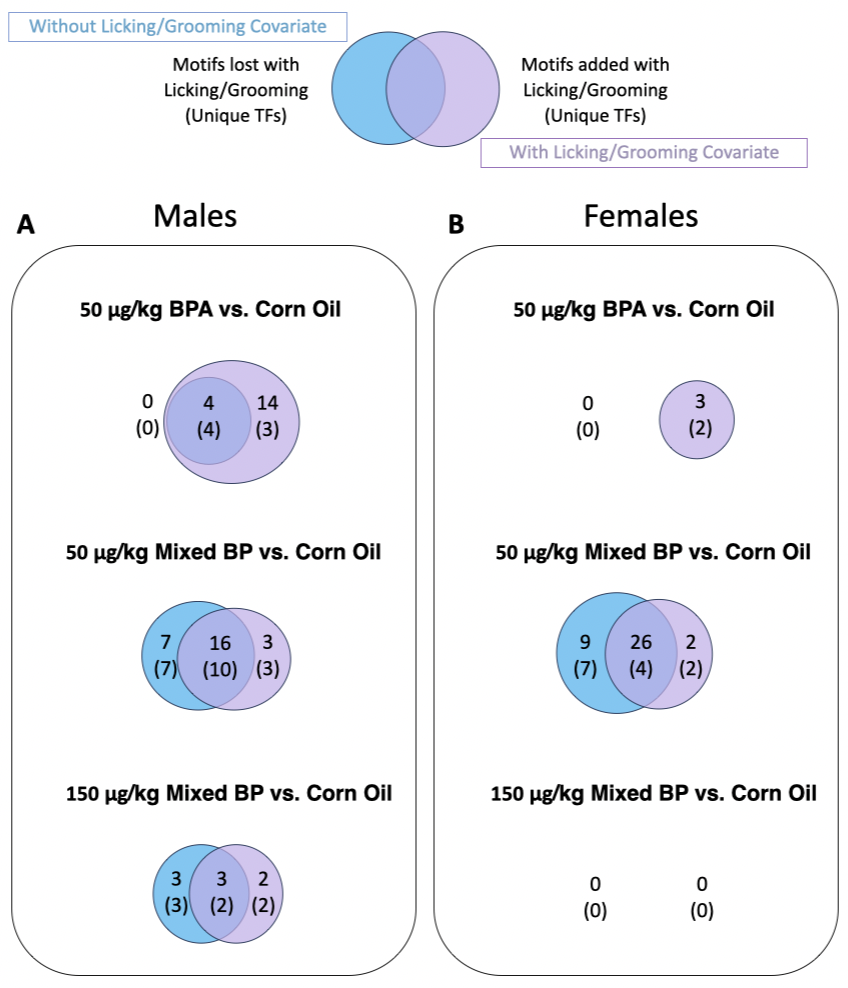

**Supplementary Fig S5.** Influence of postnatal maternal licking/grooming on differentially methylated transcription factor binding sites of prenatal bisphenol-exposed male and female pups using the SArKS method. Including maternal licking/grooming as a covariate altered overrepresented sequence motifs and predicted transcription factor binding sites in all prenatal treatment groups in male (**A**) and female (**B**) pups (except for females from the 150 μg/kg Mixed BP group). Venn diagrams are displayed with the number of motifs and unique transcription factors when excluding (blue) and including (purple) maternal licking/grooming as a covariate. Motifs that do not overlap are considered to be influenced by postnatal maternal licking/grooming.

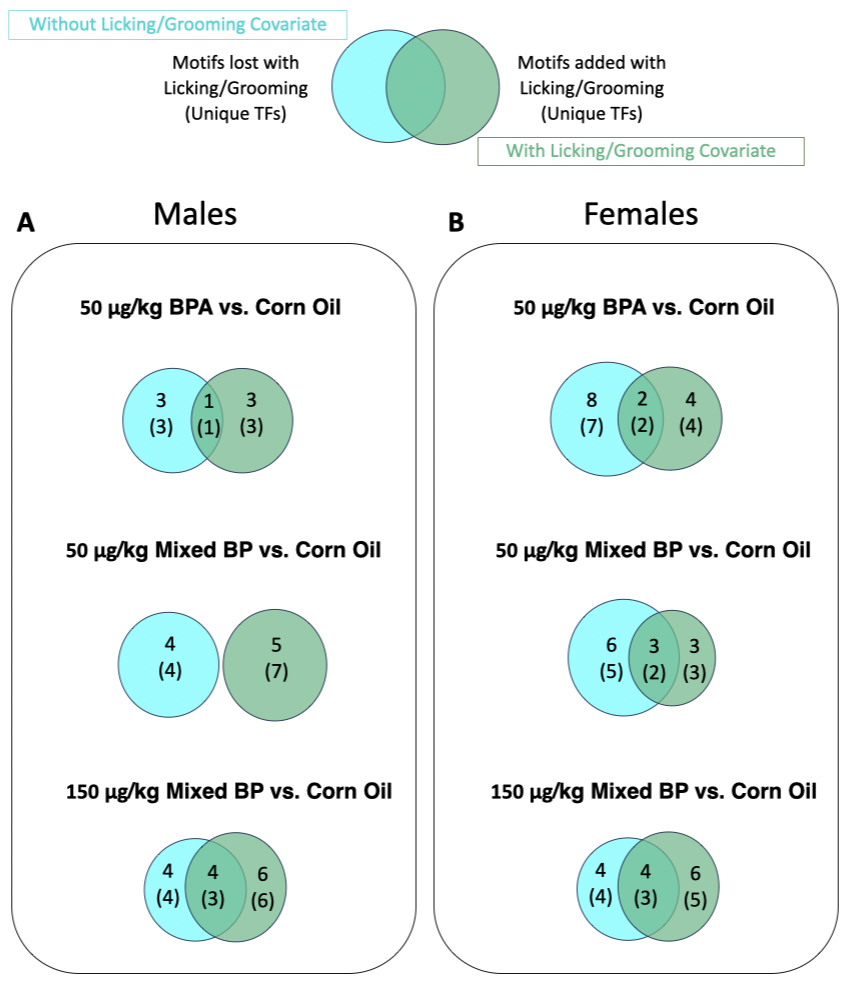

**Supplementary Fig S6.** Influence of postnatal maternal licking/grooming on differentially methylated transcription factor binding sites of prenatal bisphenol-exposed male and female pups using the MEME method. Including maternal licking/grooming as a covariate altered overrepresented sequence motifs and predicted transcription factor binding sites in most prenatal treatment groups in male (**A**) and female (**B**) pups. Venn diagrams are displayed with the number of motifs and unique transcription factors when excluding (blue) and including (green) maternal licking/grooming as a covariate. Motifs that do not overlap are considered to be influenced by postnatal maternal licking/grooming.

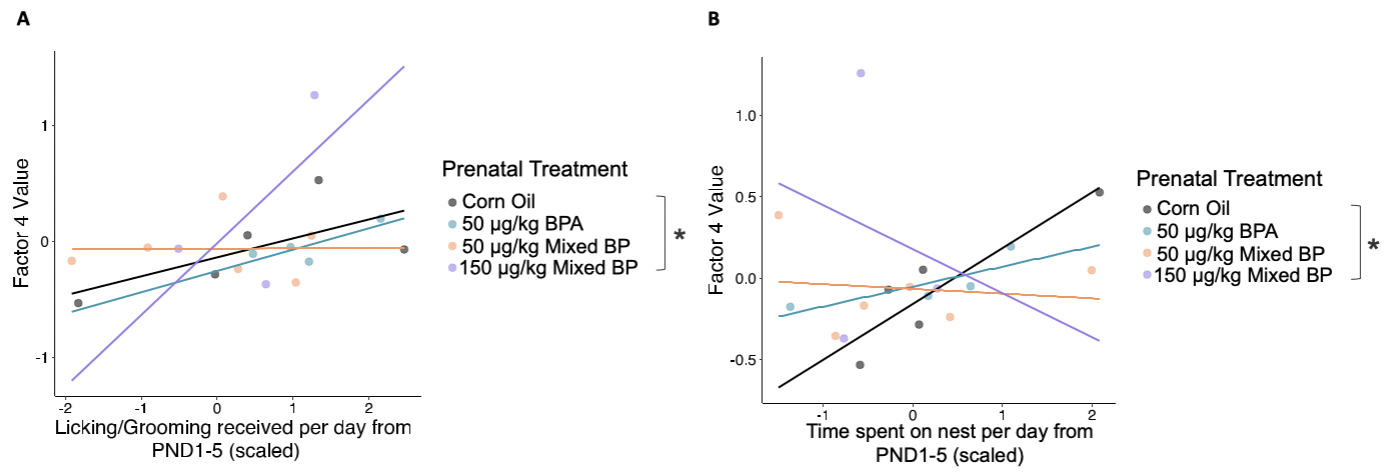

**Supplementary Fig S7.** Interactions between prenatal bisphenol exposure and postnatal maternal licking/grooming on Factor 4 generated from the multi-omic factor analysis (MOFA) in female pups. Significant interactions between the 150 μg/kg Mixed BP exposure group and maternal licking/grooming (**A**) and nest attendance (**B**) were found when combining the transcriptome and DNA methylome datasets. In females from the 150 μg/kg Mixed BP group, Factor 4 values were positively correlated with maternal licking/grooming and negatively correlated with nest attendance. Scatterplots are displayed with linear regression lines for each prenatal treatment group. * p < 0.05 interaction between prenatal treatment and postnatal maternal care
