## Supplementary Results for "Postnatal maternal care normalizes the hypothalamic DNA methylome following prenatal bisphenol exposure"

***2.3 WGCNA gene expression modules associated with interactive effects of prenatal BP and postnatal maternal LG***

The expression levels of individual genes are not independent from each other. Changes in gene expression tend to have a network-like structure of many co-expressed genes that are regulated by common transcription factors, such as estrogen receptors. Weighted Gene Co-expression Network Analysis (WGCNA) allows us to cluster these naturally occurring co-expressed gene networks into modules to examine the main effects of prenatal BP and its interactions with postnatal maternal care on differentially expressed gene networks. We can then examine if these gene modules are enriched in estrogen-responsive genes and their connectivity to estrogen and estrogen-related receptor expression. For the WGCNA analyses, a total of 7 modules for male pups and 13 modules for female pups were analyzed. For the male pups, there was only a significant main effect of prenatal treatment group found for one module, where the eigenvalues for the 50 μg/kg Mixed BP exposure group were lower than the Corn Oil group. For the female pups, there were significant or marginal interactions between the 150 μg/kg Mixed BP exposure group and maternal licking/grooming on five modules (Blue, Brown, Magenta, Turquoise, Yellow). More specifically, there were significant interactions for the Brown module (t = -2.544, p = 0.0292; 2,481 genes; **Supplementary Figure S3A**), the Turquoise module (t = 2.291, p = 0.0449; 5,229 genes; **Supplementary Figure S3B**), and the Yellow module (t = 2.061, p = 0.0264; 1,445 genes; **Supplementary Figure S3C**). In females from the 150 μg/kg Mixed BP group, maternal licking/grooming was negatively correlated with the Brown eigengene values and positively correlated with the Turquoise and Yellow eigengene values. In the Corn Oil group, the relationships between maternal licking/grooming and the WGCNA module eigengene values were reversed. There were also marginal interactions for the Blue module (t = -2.179, p = 0.0544; 4,022 genes; **Supplementary Figure S3D**) and the Magenta module (t = -2.205, p = 0.0520; 980 genes; **Supplemental Figure S3E**), Top GO terms for each of the modules with significant or marginal interactions in female offspring are presented in **Supplementary Figure S3F**. There were no significant interactions found between prenatal bisphenol exposure and nest attendance; however, including nest attendance and the interaction between prenatal bisphenol exposure and nest attendance in the linear model was required to reveal the interactions between the 150 μg/kg Mixed BP exposure group and maternal licking/grooming.

**2.6.1 DMRs influenced by maternal LG in male pups** For male pups with prenatal 50 μg/kg BPA exposure, there were 43 DMRs (corresponding to 32 genes) that were unchanged, 49 DMRs (corresponding to 31 genes) that were lost, and 54 DMRs (corresponding to 38 genes) that were added when including maternal licking/grooming as a covariate (**Supplementary Figure S4A**; **Supplementary Table S2**). In the prenatal 50 μg/kg Mixed BP group, there were 31 DMRs (corresponding to 16 genes) that were unchanged, 7 DMRs (corresponding to one gene) that were lost, and 47 DMRs (corresponding to 35 genes) that were added when including maternal licking/grooming as a covariate (**Supplementary Figure S4B**; **Supplementary Table S3**). In the prenatal 150 μg/kg Mixed BP group, there were 44 DMRs (corresponding to 27 genes) that were unchanged, 6 DMRs (corresponding to four genes) that were lost, and 38 DMRs (corresponding to 21 genes) that were added when including maternal licking/grooming as a covariate (**Supplementary Figure S4C**; **Supplementary Table S4**).

**2.6.2 DMRs influenced by maternal LG in female pups** For female pups with prenatal 50 μg/kg BPA exposure, there were 20 DMRs (corresponding to nine genes) that were unchanged, 23 DMRs (corresponding to 14 genes) that were lost, and 28 DMRs (corresponding to 15 genes) that were added when including maternal licking/grooming as a covariate (**Supplementary Figure S4D; Supplementary Table S5**). In the prenatal 50 μg/kg Mixed BP group, there were 46 DMRs (corresponding to 27 genes) that were unchanged, 25 DMRs (corresponding to 19 genes) that were lost, and three DMRs (corresponding to two genes) that were added when including maternal licking/grooming as a covariate (**Supplementary Figure S4E; Supplementary Table S6**). In the prenatal 150 μg/kg Mixed BP group, there were 60 DMRs (corresponding to 48 genes) that were unchanged, 10 DMRs (corresponding to eight genes) that were lost, and 25 DMRs (corresponding to 16 genes) that were added when including maternal licking/grooming as a covariate (**Supplementary Figure S4F; Supplementary Table S7**).

While the gene lists for each comparison were too small to run any functional enrichment analyses (e.g., GO), there were genes related to neurotransmission (e.g., Grik1, Kcnip1, Hcn2), neurodevelopment (e.g., Epha6, Nlgn3, Sash1), gene regulation (e.g., Arx, Zfp521, Mir135b), and mitochondrial function (e.g., Mtfr2, Gpam, Mthfd2) throughout all of the comparisons between prenatal bisphenol treatment and Corn Oil groups, similar to the gene modules of interest in the WGCNA analysis. There were also several notable neuroendocrine-related genes (Ddc, Maoa, Kiss1, Glpr1, Igf2bp2, Drd5, Gper1, Hsd17b2) found throughout all comparisons between prenatal bisphenol treatment and Corn Oil groups.

***2.7 Impact of postnatal maternal LG on differentially methylated transcription factor binding sites in prenatal BP-exposed pups***

To connect the changes in estrogen receptor signaling found in the tag-seq data and DNA methylation changes more directly, we next examined the influence of postnatal maternal licking/grooming on differential methylation of proximal transcription factor binding sites and EREs. To assess this outcome, we performed a motif analysis using two different methods (SArKS and MEME Suite) to identify overrepresented sequence motifs and JASPAR to match the motifs to known transcription factor binding sites.

**2.7.1 Differentially methylated transcription factor binding sites in male pups** When excluding licking/grooming as a covariate in male pups, SArKS identified four significant motifs associated with four unique transcription factors in hypomethylated regions with prenatal 50 μg/kg BPA exposure, 23 significant motifs associated with 17 unique transcription factors in hypermethylated regions with prenatal 50 μg/kg Mixed BP exposure, and six significant motifs associated with five unique transcription factors in hypomethylated regions with prenatal 150 μg/kg Mixed BP exposure (**Supplementary Table S8**). When including licking/grooming as a covariate, SArKS identified 18 significant motifs associated with seven unique transcription factors in hypomethylated regions with prenatal 50 μg/kg BPA exposure, 16 significant motifs associated with 13 unique transcription factors in hypermethylated regions with prenatal 50 μg/kg Mixed BP exposure, and four significant motifs associated with four unique transcription factors in hypomethylated regions with prenatal 150 μg/kg Mixed BP exposure (**Supplementary Table S9**). Overall, when including maternal licking/grooming as a covariate, three new transcription factor binding sites (EBF2, GLI3, TFAP2C) were identified in hypomethylated regions with prenatal 50 μg/kg BPA exposure, three new transcription factor binding sites (NRF1, E2F3, EGR3) were identified in hypermethylated regions with prenatal 50 μg/kg Mixed BP exposure, and two new transcription factor binding sites (EGR3, E2F7) were identified in hypomethylated regions with prenatal 150 μg/kg Mixed BP exposure (**Supplementary Figure S5A**).

**2.7.2 Differentially methylated transcription factor binding sites in female pups** When excluding licking/grooming as a covariate in female pups, SArKS identified 34 significant motifs associated with 10 unique transcription factors in hypomethylated regions and two significant motifs associated with two transcription factors in hypermethylated regions with prenatal 50 μg/kg Mixed BP exposure only (**Supplementary Table S10**). When including licking/grooming as a covariate, SArKS identified three significant motifs associated with two unique transcription factors in hypermethylated regions with prenatal 50 μg/kg BPA exposure as well as two significant motifs associated with two unique transcription factors in hypomethylated regions and four significant motifs associated with four transcription factors in hypermethylated regions with prenatal 50 μg/kg Mixed BP exposure (**Supplementary Table S11**). Overall, when including maternal licking/grooming as a covariate, two new transcription factor binding sites (SP5, ZNF610) were identified with prenatal 50 μg/kg BPA exposure and two new transcription factor binding sites (HAND2, IKZF2) were identified with prenatal 50 μg/kg Mixed BP exposure, all within hypermethylated regions (**Supplementary Figure S5B**).

There were several predicted transcription factor binding sites that reappeared using the MEME Suite method in male offspring (SP5, HAND2, RBPJ) and female offspring (ELK1, IKZF2, SP5, RBPJ, HAND2). There were also changes in motifs and predicted transcription factor binding sites when adding postnatal maternal licking/grooming as a covariate with MEME Suite (**Supplementary Figure S6**). In many comparisons, there were a larger number of motifs and predicted transcription factor binding sites that were altered with postnatal maternal licking/grooming and less overlap using the MEME Suite method than the SArKS method.

While no estrogen receptors were found in the list of predicted transcription factor binding sites, many of the predicted transcription factors have known interactions with estrogen receptors. This includes transcription factors that are known to dimerize with ESR1, block estrogen receptors from binding to DNA, facilitate transcription of *Esr1*, or are estrogen-responsive genes themselves (**Supplementary Table S12**).

***2.8 Interactive effects of prenatal BP exposure and postnatal maternal LG on the multi-omic epigenome in the MPOA of female pups***

The disconnect between the gene expression and DNA methylation changes may be in part due to the different analysis methods used to examine the interactions between prenatal BP exposure and maternal care. To address this disparity, we also examined the influence of postnatal maternal licking/grooming on the combined DNA methylome and transcriptome in the developing MPOA using Multi-Omic Factor Analysis (MOFA). For male pups, there were two factors that showed significant interactions of prenatal bisphenol exposure and maternal licking/grooming; however, upon further inspection these interactions appear to be solely driven by one outlier in both factors (data not shown). For female pups, there was one factor (Factor 4) that showed significant interactions between the 150 μg/kg Mixed BP exposure group and maternal licking/grooming (t = 4.456, p = 0.0043; **Supplementary Figure S7A**) as well as nest attendance (t = 3.182, p = 0.0190; **Supplementary Figure S7B**). Factor 4 values were more strongly positively correlated with maternal licking/grooming in females from the 150 μg/kg Mixed BP group than the Corn Oil group. Factor 4 values were negatively correlated with nest attendance in the 150 μg/kg Mixed BP group, though the smaller range of nest attendance values limit full interpretation of the data. In the Corn Oil group, Factor 4 values were positively correlated with nest attendance. Similar to the WGCNA analysis, including nest attendance and the interaction between prenatal bisphenol exposure and nest attendance in the linear model was required to reveal the significant interaction between the 150 μg/kg Mixed BP exposure group and maternal licking/grooming.
